# Metabolism of long-chain fatty acids affects disulfide bond formation in *Escherichia coli* and activates envelope stress response pathways as a combat strategy

**DOI:** 10.1101/2020.03.04.976175

**Authors:** Kanchan Jaswal, Megha Shrivastava, Deeptodeep Roy, Shashank Agrawal, Rachna Chaba

**Affiliations:** Department of Biological Sciences, Indian Institute of Science Education and Research (IISER) Mohali, SAS Nagar, Punjab, India

## Abstract

The envelope of gram-negative bacteria serves as the first line of defense against environmental insults. Therefore, its integrity is continuously monitored and maintained by several envelope stress response (ESR) systems. Due to its oxidizing environment, the envelope represents an important site for disulfide bond formation. In *Escherichia coli*, the periplasmic oxidoreductase, DsbA introduces disulfide bonds in substrate proteins and transfers electrons to the inner membrane oxidoreductase, DsbB. Under aerobic conditions, the reduced form of DsbB is re-oxidized by ubiquinone, an electron carrier in the electron transport chain (ETC). Given the critical role of ubiquinone in transferring electrons derived from the oxidation of reduced cofactors, we were intrigued whether metabolic conditions that generate a large number of reduced cofactors render ubiquinone unavailable for disulfide bond formation. To test this, here we investigated the influence of metabolism of long-chain fatty acid (LCFA), an energy-rich carbon source, on the redox state of the envelope. We show that LCFA degradation increases electron flow in the ETC. Further, we find that whereas cells metabolizing LCFAs exhibit several characteristics of insufficient disulfide bond formation, these hallmarks are averted in cells exogenously provided with ubiquinone. Importantly, the ESR pathways, Cpx and σ^E^, are activated by envelope signals generated during LCFA metabolism, and these systems maintain proper disulfide bond formation. We find that σ^E^ downregulation hampers disulfide bond formation only in the absence of Cpx, and amongst the two ESR systems, only Cpx senses redox-dependent signal and is induced to a greater extent by LCFAs. Therefore, we argue that Cpx is the primary ESR that senses and maintains envelope redox homeostasis. Taken together, our results demonstrate an intricate relationship between cellular metabolism and disulfide bond formation dictated by ETC and ESR, and provide the basis for examining whether similar mechanisms control envelope redox status in other gram-negative bacteria.

**Author summary:** Disulfide bonds contribute to the folding and stability of many extracytoplasmic proteins in all domains of life. In gram-negative bacteria, including *Escherichia coli*, disulfide bond formation occurs in the oxidizing environment of the periplasmic space enclosed within the outer and inner membrane layers of the envelope. Because disulfide-bonded proteins are involved in diverse biological processes, bacteria must monitor the envelope redox status and elicit an appropriate response when perturbations occur; however, these mechanisms are not well elucidated. Here, we demonstrated that the metabolism of an energy-rich carbon source, long-chain fatty acid (LCFA) hampers disulfide bond formation in *E. coli*. An envelope stress response (ESR) system, Cpx, senses this redox imbalance and maintains proper disulfide bond formation. The σ^E^ pathway, another ESR system, plays an ancillary role in maintaining redox homeostasis. LCFA metabolism, disulfide bond formation, and ESR systems have independently been implicated in the pathogenesis of several gram-negative bacteria. The present study sets the basis to explore whether LCFA metabolism impacts the virulence of these bacteria by influencing the redox status of their envelope and activation of ESR pathways.

## Introduction

The multilayered envelope of gram-negative bacteria comprised of the inner membrane, the outer membrane and the peptidoglycan layer within the periplasm, protects the cell from environmental stresses and is the site for a myriad of functions critical for cellular growth and viability (reviewed in [1]). Given the essential role of the envelope in maintaining cellular homeostasis, its integrity must be continuously monitored. In *E. coli*, this task is accomplished by at least five dedicated envelope stress response (ESR) systems (Bae, Cpx, Psp, Rcs and σ^E^), which sense problems in the envelope and change the transcriptional program to combat stress (reviewed in [2, 3]). For example, the Cpx pathway predominantly deals with misfolded inner membrane and periplasmic proteins, and defects in inner membrane protein translocation, peptidoglycan biosynthesis and lipoprotein trafficking, while the σ^E^ pathway senses and responds to defects in the transport and assembly of outer membrane proteins (OMPs) and lipopolysaccharide (LPS) [4-6].

A critical function of the envelope of gram-negative bacteria is oxidative protein folding. Many secreted proteins require disulfide bonds for their maturation and stability. In *E. coli*, ∼300 proteins are predicted to be disulfide-bonded in the periplasm. The formation of disulfide bonds in *E. coli* takes place with the help of a periplasmic oxidoreductase, DsbA, which gets reduced after catalyzing the formation of disulfide bonds in substrate proteins. DsbB, an inner membrane disulfide oxidoreductase, performs the re-oxidation of DsbA [7, 8]. Due to the requirement of disulfide bond formation in proteins involved in diverse biological processes, mutants of disulfide bond-forming machinery show pleiotropic phenotypes. For example, *dsbA* and *dsbB* mutants of *E. coli* exhibit dramatic reductions in motility and alkaline phosphatase (AP) activity, and sensitivity to thiol agents such as DTT, heavy metals such as cadmium and zinc, and drugs such as benzylpenicillin [9-14]. Furthermore, mutants of several pathogenic bacteria defective in disulfide bond formation are attenuated for virulence [15].

The ultimate oxidizing power for disulfide bond formation is provided by the electron transport chain (ETC). The reduced form of DsbB is re-oxidized by transferring electrons to the quinones, ubiquinone and menaquinone, under aerobic and anaerobic conditions, respectively. The terminal oxidases finally shuttle electrons from reduced quinones (quinol) to the terminal electron acceptors, for example, cytochrome *bo* transfers electrons from reduced ubiquinone (ubiquinol) to molecular oxygen under aerobic conditions. This link between ETC and envelope redox homeostasis was demonstrated in earlier *in vivo* studies where the status of disulfide bond formation was examined under conditions of non-operational or defective ETC. For example, *E. coli* grown in a purely fermentative manner, without any terminal electron acceptor, i.e., non-operational ETC, was compromised for disulfide bond formation, and mutants defective in the biosynthesis of the respiratory chain components, heme and quinones accumulated DsbA in a reduced form [16, 17].

In the presence of terminal electron acceptors, i.e., operational ETC, besides transferring electrons from the disulfide bond-forming machinery, quinones play a pivotal role in transferring electrons generated by the metabolism of carbon sources from respiratory dehydrogenases to the terminal oxidases [18]. The convergence of metabolism and disulfide bond formation at the level of quinones in the ETC makes it tempting to speculate that metabolic conditions that increase electron flow in the ETC might limit quinones from taking up electrons from the disulfide bond-forming machinery, thereby compromising envelope redox homeostasis. So far, this scenario has not been investigated. Besides a fundamental perspective, because disulfide bonds are required for the activity of several virulence factors, these studies are of tremendous importance to envision how carbon metabolism impacts bacterial pathogenesis.

Long-chain fatty acids (LCFAs), carboxylic acids with a linear aliphatic chain of 12-20 carbon atoms, are used as a rich source of metabolic energy by *E. coli* and several other bacterial pathogens [19-21]. The metabolic pathway of LCFAs suggests that its degradation generates a large number of reduced cofactors, which might increase electron flow in the ETC [22]. Thus, LCFAs represent a suitable carbon source to examine the interconnection between cellular metabolism and the redox status of the envelope.

Here, we investigated the relation between LCFA metabolism and disulfide bond formation in *E. coli* K-12. We show that LCFA metabolism increases electron flow in the ETC. We find that LCFA-utilizing cells exhibit hallmarks of insufficient disulfide bond formation, and these are prevented in cells exogenously provided with ubiquinone. Collectively, these data establish that during LCFA metabolism, ubiquinone is limiting for disulfide bond formation. We show that the envelope signals generated during growth in LCFAs activate the ESR systems, Cpx and σ^E^, which maintain proper disulfide bond formation. Our results suggest that whereas Cpx is the primary response that maintains envelope redox homeostasis, the σ^E^ pathway plays an auxiliary role in this process. Taken together, the present study emphasizes that ETC and ESR systems govern the interconnection between cellular metabolism and envelope redox homeostasis.

## Results

### LCFA metabolism increases electron flow in the ETC

Based on the metabolic pathway, LCFA degradation theoretically generates a large number of reduced cofactors. One round of β-oxidation produces one molecule each of NADH and FADH_2_ and releases two carbon atoms as acetyl-CoA. The metabolism of acetyl-CoA in the tricarboxylic acid (TCA) cycle further generates two NADH and one FADH_2_ (Fig 1A) [22]. Thus during degradation of oleate (C18:1 cis-9), a representative LCFA used in this study, β-oxidation will produce eight molecules each of NADH and FADH_2_, and the TCA cycle will further generate more reduced cofactors from nine molecules of acetyl-CoA. We tested whether oleate catabolism indeed produces a large number of reduced cofactors. For this, WT BW25113 was grown in buffered tryptone broth (TBK) supplemented either with oleate (TBK-Ole) or with the detergent Brij-58 used for solubilizing oleate (TBK-Brij) (Fig 1B), and steady-state levels of NADH and NAD^+^ were measured in exponential phase (time point T1, Fig 1B). These and subsequent experiments were carried out in TBK-Ole to support the growth of control *fad* knockouts, which otherwise do not grow in minimal medium containing oleate as a sole carbon source [23]. Moreover, because tryptone causes mild catabolite repression [24], oleate is co-utilized with carbon components of tryptone [23]. We observed ∼2.5-fold increase in NADH in WT cells grown in TBK-Ole compared to TBK-Brij; however, there was no significant change in NAD^+^ levels. Overall, this resulted in ∼2.5-fold higher NADH/NAD^+^ ratio in TBK-Ole (0.363 ± 0.017) compared to TBK-Brij (0.135 ± 0.007). The higher NADH level and NADH/NAD^+^ ratio were due to oleate utilization; in contrast to WT, a Δ*fadL* strain (*fadL* codes for the outer membrane transporter for LCFAs) had similar NADH level and NADH/NAD^+^ ratio in TBK-Ole and TBK-Brij (Fig 1C).

**Figure 1.**
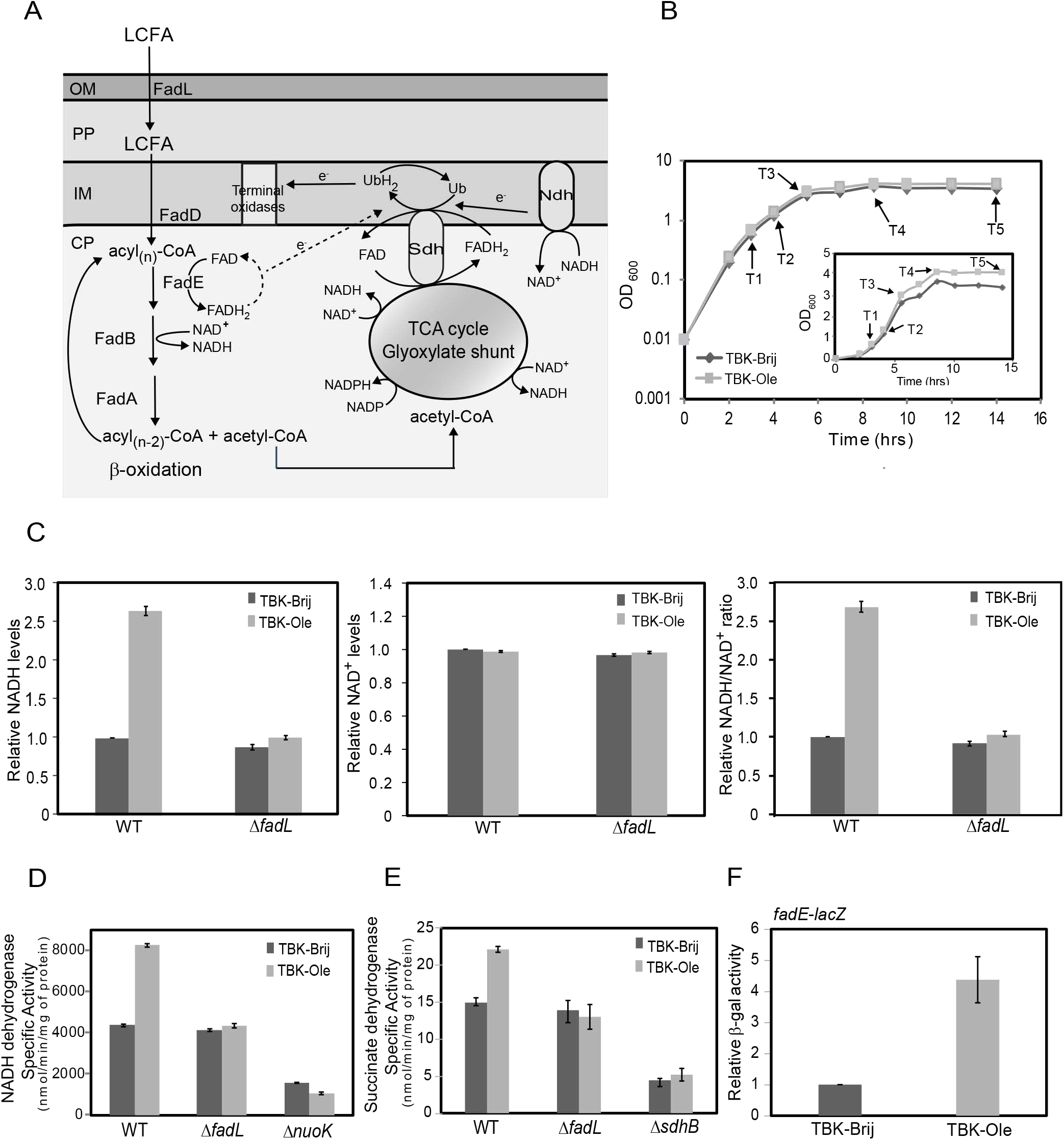
LCFA metabolism increases electron flow in the ETC. *A)* Pathway of LCFA transport and degradation in *E. coli*. FadL transports exogenously supplied LCFAs across the outer membrane. LCFAs are extracted from the inner membrane by the acyl-CoA synthetase, FadD, which also esterifies LCFAs to acyl-CoA. Acyl-CoAs are degraded to acetyl-CoA via β-oxidation mediated by the enzymes FadE, FadB, and FadA. Acetyl-CoA is further metabolized in the TCA and glyoxylate cycles. NADH and FADH_2_ produced during β-oxidation and TCA cycle are oxidized in the ETC by respiratory dehydrogenases. The electrons are transferred to the lipid-soluble electron carrier, ubiquinone. The reduced form of ubiquinone, ubiquinol, further donates electrons to the terminal oxidases. Arrows with *e*^-^ labeled on the line denote the direction of electron flow. The dotted arrow indicates that the components involved in the oxidation of FadE and electron transfer from FadE to the ETC are not established. Abbreviations: CP, cytoplasm; IM, inner membrane; PP, periplasm; OM, outer membrane; TCA, tricarboxylic acid; Ndh, NADH dehydrogenases; Sdh, succinate dehydrogenase; Ub, ubiquinone; UbH_2_, ubiquinol. *B) Growth curve of WT BW25113 in TBK-Brij and TBK-Ole.* WT was grown either in TBK-Brij or TBK-Ole. OD_600_ of the cultures was measured, and growth curves were plotted on a semi-logarithmic scale. The experiment was done three times. A representative dataset is shown. T1, T2, T3, T4, and T5 indicate time points where cultures were harvested for various assays. *Inset:* The above growth curves were also plotted on a linear scale. *C) NADH/NAD*^*+*^ *ratio is higher in oleate-utilizing cells.* Strains were grown either in TBK-Brij or TBK-Ole. Cultures were harvested at time point T1 indicated in Fig 1B, and NADH (left panel) and NAD^+^ (middle panel) levels, and NADH/NAD^+^ ratio (right panel) were determined. Data were normalized to WT in TBK-Brij and represent average (± S.D.) of three independent experiments. *D and E) The activity of NADH dehydrogenase (D) and succinate dehydrogenase (E) increases in cells utilizing oleate.* Strains were grown either in TBK-Brij or TBK-Ole. Cultures were harvested at time point T1 indicated in Fig 1B, and the activity of the respiratory dehydrogenases was measured. Data represent the average (± S.D.) of three independent experiments. *F) FadE is induced in oleate-utilizing cells.* WT carrying chromosomal fusion of *lacZ* with the promoter of *fadE* was grown either in TBK-Brij or TBK-Ole. Cultures were harvested at time point T1 indicated in Fig 1B, and β-gal activity was measured. Data were normalized to the β-gal activity of WT in TBK-Brij. Data represent the average (± S.D.) of four independent experiments. The average β-gal activity of the reporter strain in TBK-Brij was 54 (± 9) Miller units.

During aerobic metabolism, NADH and FADH_2_ are oxidized at the ETC by NADH dehydrogenases and succinate dehydrogenase, respectively, and the electrons are transferred to the lipid-soluble electron carrier, ubiquinone (Fig 1A) [18]. We investigated whether a large amount of reduced cofactors produced during LCFA metabolism increases electron flow in the ETC by determining the activity of NADH and succinate dehydrogenases. The activity of both the respiratory dehydrogenases was higher in WT cultured in TBK-Ole compared to cells grown in TBK-Brij and this increase was dependent on oleate utilization; the increase in activity was abolished in a Δ*fadL* strain. As expected, the activity of NADH and succinate dehydrogenases was considerably lower in Δ*nuoK* (*nuoK* codes for a subunit of NADH dehydrogenase I) and Δ*sdhB* (*sdhB* codes for a subunit of succinate dehydrogenase) strains, respectively, in both TBK-Brij and TBK-Ole (Figs 1D and E). We previously reported an increase in transcript levels of an acyl-CoA dehydrogenase, FadE, in *E. coli* cultured in TB medium supplemented with oleate (compared to TB-Brij control) [23]. FadE catalyzes the oxidation of acyl-CoA (an intermediate in LCFA metabolism) concomitant with reduction of FAD to FADH_2_ and has also been speculated to re-oxidize FADH_2_ to FAD by transferring electrons from its dehydrogenase domain to the ETC (Fig 1A) [25]. Since here we used buffered TBK media for all our experiments, we checked the induction of *fadE* in TBK-Ole at the exponential phase by assaying β-galactosidase (β-gal) activity of chromosomal *lacZ* fused to the promoter of *fadE*. We observed ∼4 fold induction of *fadE* in cells grown in TBK-Ole in comparison to TBK-Brij (Fig 1F). This data is consistent with our previous observation of increased *fadE* transcript levels in oleate [23] and collectively suggests an increased flow of electrons from FadE to the ETC during LCFA metabolism.

### Ubiquinone is limiting for disulfide bond formation during LCFA metabolism

Because electrons from aerobic metabolism of carbon sources and disulfide bond-forming machinery converge at the level of ubiquinone in the ETC, we suggested that a large flow of electrons during oleate metabolism would make ubiquinone less available to handle electrons from the disulfide bond-forming machinery. To test this proposal, we investigated whether oleate-grown cells show characteristics of insufficient disulfide bond formation.

Alkaline phosphatase (AP) is a periplasmic enzyme that requires intramolecular disulfide bonds for its activity [26]; hence AP activity is routinely used to determine the redox state of the periplasm [27]. As a test for our suggestion that LCFA metabolism results in a less oxidizing environment in the periplasm, we cultured RI89 strain (a *phoR* mutant that constitutively expresses AP) [27], in TBK-Ole and TBK-Brij (Fig 2A), and measured AP activity in the exponential phase. In line with our hypothesis, compared to TBK-Brij, in TBK-Ole, there was ∼40% reduction in AP activity (Fig 2B). This decrease was not due to the difference in AP protein levels (*Inset* Fig 2B). The reduction in AP activity was due to oleate utilization; compared to RI89, the AP activity did not exhibit any decrease in TBK-Ole in the isogenic Δ*fadL* and Δ*fadE* strains (Fig 2B). Furthermore, although in TBK-Brij, the AP activity decreased to ∼30% and ∼60% in Δ*dsbA* and Δ*dsbB* strains, respectively, the activity did not drop further in TBK-Ole (Fig 2C). These data validate that insufficient disulfide bond formation in the periplasmic enzyme in TBK-Ole is due to defects in the DsbA-DsbB oxidative pathway. Importantly, exogenous supplementation of ubiquinone-8 (the form of ubiquinone naturally present in the ETC of *E. coli* [28]) prevented a decrease in AP activity of RI89 grown in TBK-Ole (Fig 2D). This result emphasizes that ubiquinone is indeed limiting for disulfide bond formation in oleate-utilizing cells.

**Figure 2.**
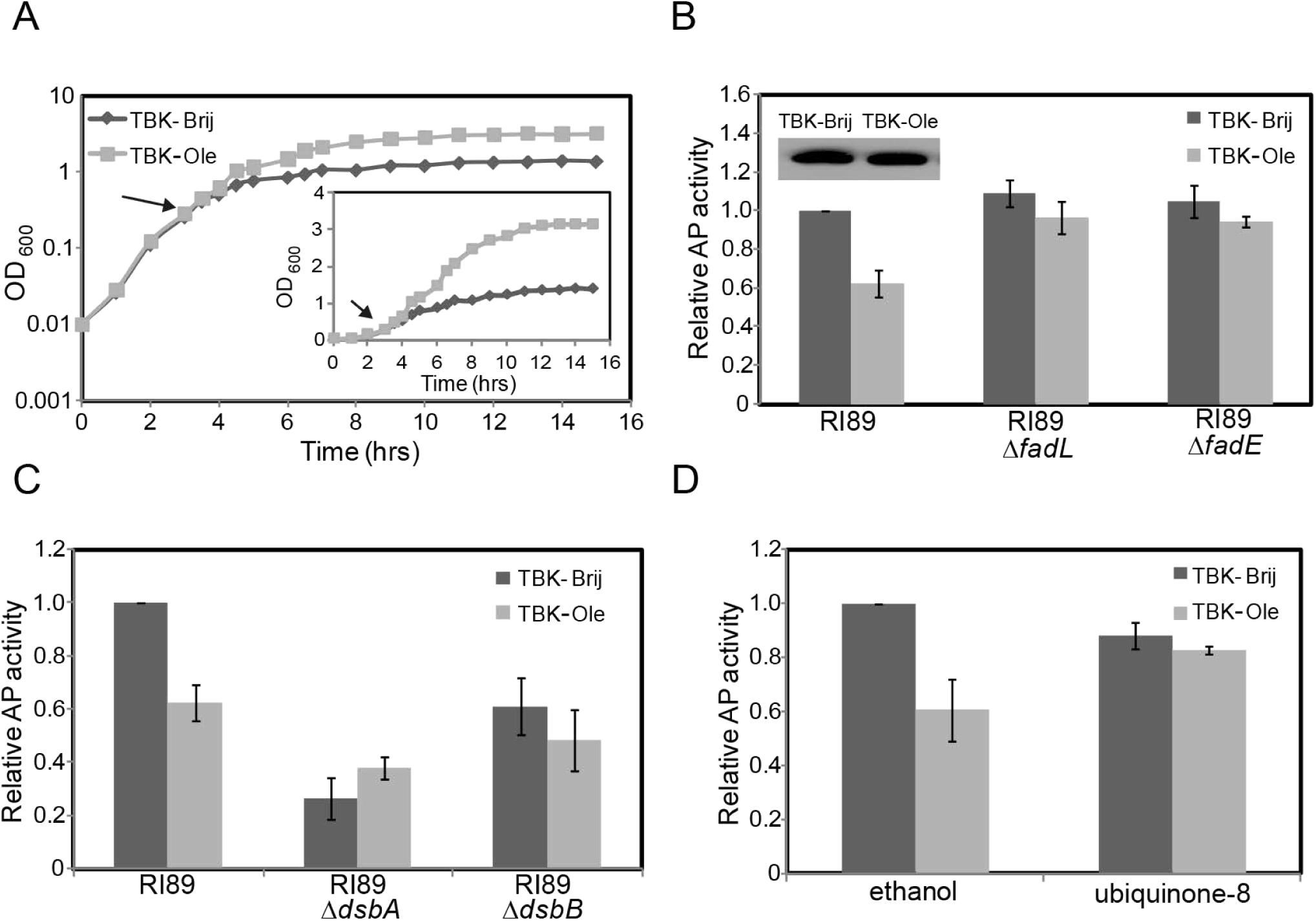
Insufficient disulfide bond formation results in a decrease in alkaline phosphatase activity in LCFA-utilizing cells. *A) Growth curve of RI89 in TBK-Brij and TBK-Ole.* RI89 was grown either in TBK-Brij or TBK-Ole. OD_600_ of the cultures was measured, and growth curves were plotted on a semi-logarithmic scale. The experiment was done three times. A representative dataset is shown. Arrow indicates the time point where the cultures were harvested for determining AP activity and AP levels. *Inset:* The above growth curves were also plotted on a linear scale. *B) Oleate utilization results in a decrease in alkaline phosphatase activity.* Cultures grown either in TBK-Brij or TBK-Ole were harvested in the exponential phase and processed for AP assay. Data were normalized to the AP activity of RI89 in TBK-Brij and represent average (± S.D.) of three independent experiments. The average AP activity of RI89 in TBK-Brij was 1384 (± 73) units. *Inset: AP levels are similar in RI89 grown in TBK-Brij and TBK-Ole.* Cultures were harvested, lysates were prepared, samples were run on 15% SDS-PAGE, and processed for Western blotting using an anti-AP antibody. The band corresponding to AP is shown (Mol. wt. ∼52 kDa). *C) Compromised DsbA-DsbB machinery is responsible for the decrease in alkaline phosphatase activity in cells utilizing oleate.* Cultures grown either in TBK-Brij or TBK-Ole were harvested in the exponential phase and processed for AP assay. Data were normalized to the AP activity of RI89 in TBK-Brij and represent average (± S.D.) of three independent experiments. The average AP activity of RI89 in TBK-Brij was 1245 (± 357) units. *D) Supplementation of ubiquinone-8 prevents a decrease in AP activity in oleate-utilizing cells.* RI89 cells were grown either in TBK-Brij or TBK-Ole. Media contained either 20 μM ubiquinone-8 or 0.1% ethanol (solvent for ubiquinone-8). Cultures were harvested in the exponential phase, AP activity was measured and the data were normalized to the AP activity of RI89 in TBK-Brij containing 0.1% ethanol. Data represent the average (± S.D.) of three independent experiments. The average AP activity of RI89 in TBK-Brij supplemented with 0.1% ethanol was 1192 (± 341) units.

Another hallmark of impaired disulfide bond formation is the hypersensitivity to thiol agents. For example, the *dsbA* and *dsbB* mutants show growth defect/ no growth on LB agar containing millimolar concentrations of dithiothreitol (DTT) that otherwise do not affect the growth of WT strain [10, 29, 30], and their growth in liquid minimal medium (glucose as carbon source) is also inhibited by millimolar concentrations of 1-thioglycerol [31]. Further, the *ubi* mutants that contain low levels of ubiquinone are also compromised for growth in minimal medium (glucose as carbon source) supplemented either with DTT or 1-thioglycerol, due to the inability of the suboptimal respiratory chain to re-oxidize Dsb enzymes reduced in the presence of excess thiol agents [31]. To test whether cells grown in oleate are also sensitive to thiols, we compared the growth of WT BW25113 strain on TBK-Brij and TBK-Ole agar containing various concentrations of DTT. Whereas growth was observed in TBK-Brij even at 8 mM DTT, in TBK-Ole, 7 mM DTT resulted in a considerable growth defect and almost no growth was observed in 8 mM DTT (Fig 3A). We also profiled the growth of WT strain in liquid minimal medium containing oleate as a sole carbon source in the presence of either DTT or 1-thioglycerol and compared it with growth in minimal medium containing acetate. We chose acetate for comparison because *i)* acetate and oleate follow a common metabolic route; they are both non-fermentable carbon sources that directly enter the TCA cycle after conversion to acetyl-CoA, and *ii)* acetate metabolism theoretically generates less amount of reduced cofactors than oleate; hence the load of electrons on ETC from metabolism would be less during growth in this carbon source [22]. Consistent with our proposal, whereas 0.5 mM DTT resulted in a substantial delay in the growth of cells in oleate, it did not have an adverse effect on growth in acetate. Similarly, whereas 1 mM 1-thioglycerol completely inhibited growth in oleate, it only had a mild effect on growth in acetate (Fig 3B).

**Figure 3.**
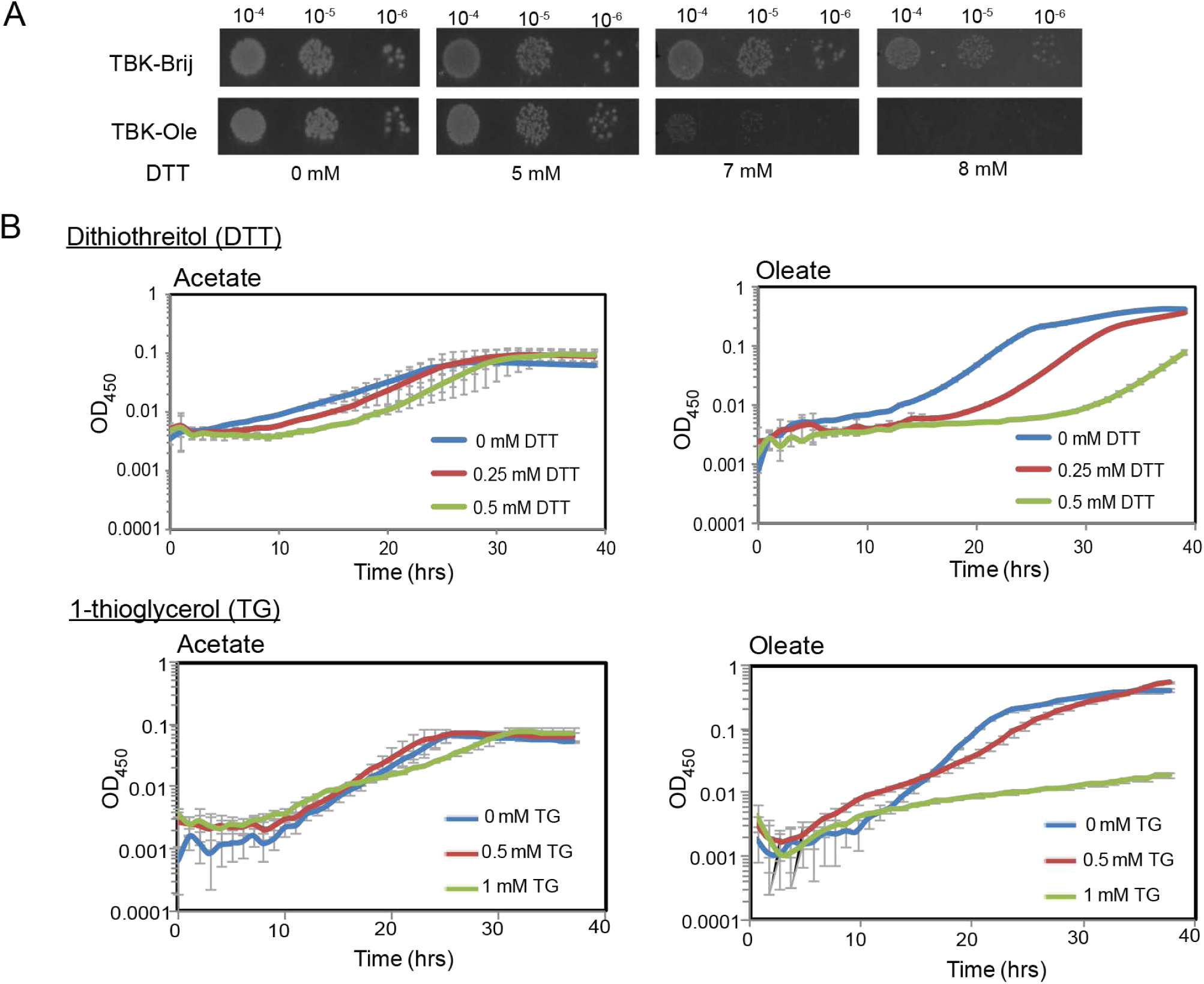
LCFA-utilizing cells exhibit thiol hypersensitivity. *A) WT cells grown in TBK-Ole are hypersensitive to DTT.* WT cells were spotted on TBK-Brij or TBK-Ole with or without indicated concentrations of DTT. The experiment was performed two times. A representative dataset is shown. *B) Amongst acetate and oleate, oleate utilization renders cells more sensitive to thiol agents.* WT cells were grown in minimal medium containing either acetate or oleate, with or without indicated concentrations of DTT or 1-thioglycerol. Acetate medium was also supplemented with Brij-58. OD_450_ of the cultures was measured, and growth curves were plotted on a semi-logarithmic scale. The experiment was done at least two times; each experiment had three technical replicates. A representative dataset, with average (± S.D.) of technical replicates, is shown.

Because of the obstruction of electron flow from the DsbA-DsbB machinery to the ETC, strains defective in quinone biosynthesis accumulate DsbA in a reduced form [17]. We reasoned that if there is limited availability of ubiquinone for disulfide bond formation during oleate metabolism, then cells might accumulate reduced form of DsbA. To test this, we harvested WT BW25113 grown in TBK-Brij and TBK-Ole at different phases of growth (Fig 1B), denatured and precipitated proteins, and treated samples with 4-acetamido-4’-maleimidystilbene-2,2’-disulfonicacid (AMS), a maleimide derivative that results in a specific, rapid and irreversible alkylation of free thiols adding ∼0.5 kDa moiety per thiol group [17]. AMS modification enabled differentiation of the oxidized and reduced forms of the thiol-containing DsbA protein on non-reducing SDS-PAGE gels. Whereas DsbA was present in the oxidized form in TBK-Brij at all phases of growth, TBK-Ole grown cells accumulated DsbA in a completely reduced form at time point T3 (corresponding to entry into stationary phase) (Figs 1B and 4A). Importantly, DsbA was present in its oxidized form at time point T3 when the TBK-Ole medium was supplemented with ubiquinone-8, reiterating that ubiquinone is limiting in oleate-utilizing cells (Fig 4B). We noted that whereas AP activity was compromised in RI89 in TBK-Ole even in the exponential phase, DsbA was completely oxidized in BW25113 in this phase of growth (compare Figs 2B and 4A). We suggest that this difference is due to the constitutive expression of AP in RI89 that likely overburdens the DsbA-DsbB machinery in TBK-Ole from the initial stages of growth.

**Figure 4.**
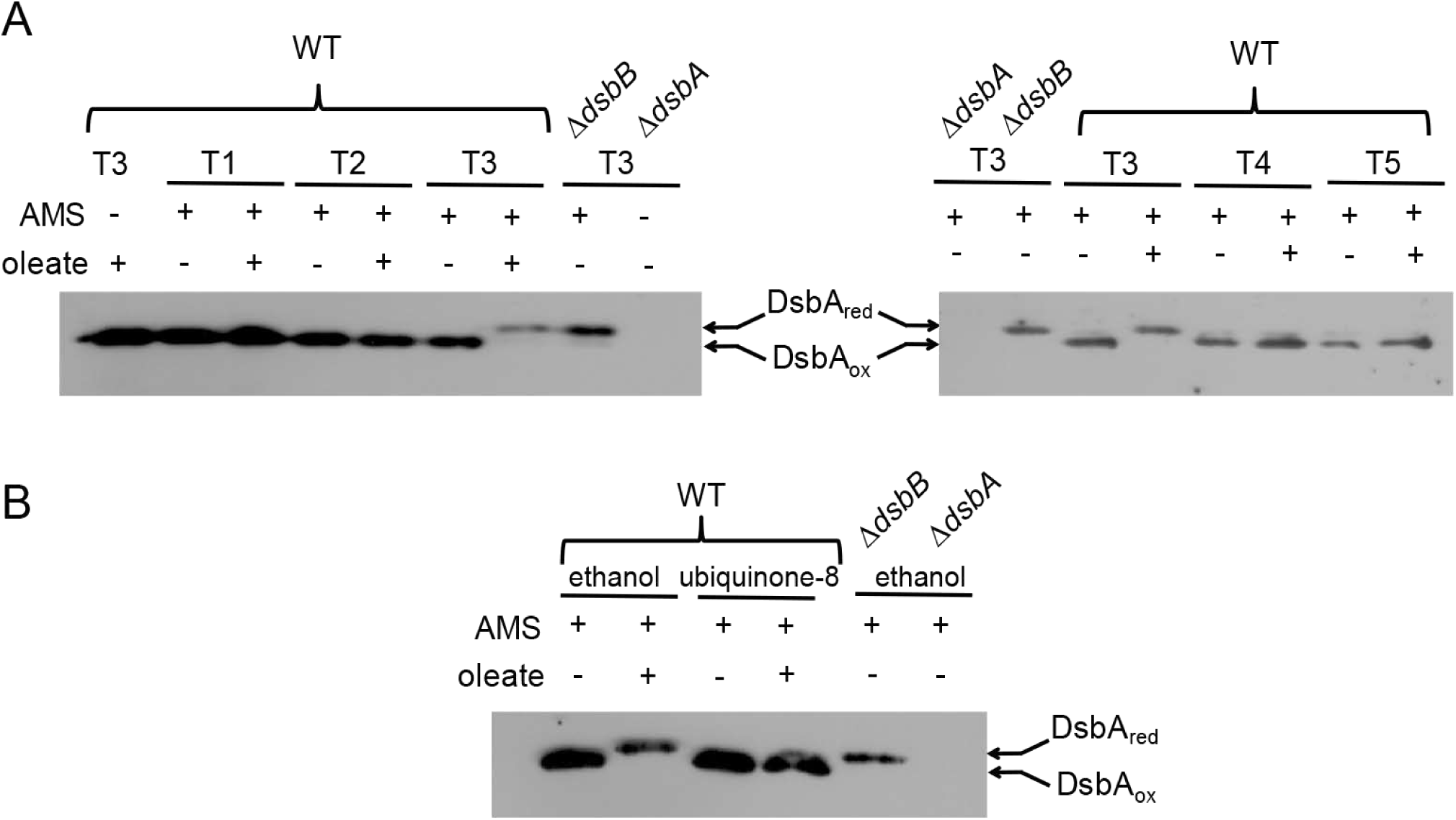
Ubiquinone limitation results in the accumulation of DsbA in a reduced form in LCFA-utilizing cells. *A) DsbA accumulates in a reduced form in oleate-utilizing cells.* WT BW25113 was grown either in TBK-Brij or TBK-Ole, and cultures were harvested at different time points, as indicated in Fig 1B. Proteins were denatured and precipitated using trichloroacetic acid, followed by treatment with AMS. Oxidized and reduced forms of DsbA were identified by running the AMS-treated samples on non-reducing SDS-PAGE gels, followed by Western blot using an anti-DsbA antibody. Δ*dsbA* and Δ*dsbB* cultured in TBK-Brij served as controls. DsbA is re-oxidized by DsbB; hence DsbA accumulates in a reduced form in a Δ*dsbB* strain. Arrows indicate oxidized and reduced forms of DsbA. *B) Supplementation of ubiquinone-8 prevents the accumulation of reduced form of DsbA in oleate-utilizing cells.* WT BW25113 and its isogenic Δ*dsbA* and Δ*dsbB* strains were grown either in TBK-Brij or TBK-Ole. The media contained either 20 μM ubiquinone-8 or 0.1% ethanol. Cultures were harvested at time point T3 (indicated in Fig 1B) and processed as mentioned in the legend to Fig 4A. Δ*dsbA* and Δ*dsbB* cultured in TBK-Brij containing 0.1% ethanol served as controls. Arrows indicate oxidized and reduced forms of DsbA.

The Δ*dsbA* strain shows sensitivity to heavy metals such as cadmium (Cd^2+^) and zinc (Zn^2+^), due to the binding of these metals with free thiols of proteins [12, 13]. Since DsbA accumulates in a reduced state in TBK-Ole, we checked if TBK-Ole-grown cells are more sensitive to Cd^2+^ compared to TBK-Brij. For this, different concentrations of cadmium chloride (CdCl_2_) were added to cultures at time point T3, and growth was monitored. Whereas there was no effect of even 2 mM CdCl_2_ on the growth of WT in TBK-Brij, TBK-Ole grown WT cells were sensitive to CdCl_2_ similar to the Δ*dsbA* strain grown in TBK-Brij (S1 Fig).

Taken together, the phenotypes of decreased AP activity, thiol hypersensitivity, accumulation of reduced form of DsbA, and cadmium sensitivity exhibited by oleate-utilizing cells convincingly establish that ubiquinone is limiting for disulfide bond formation during growth in LCFAs.

### Cpx and σ^E^ pathways are activated in LCFA-utilizing cells

In our experiment above, where we monitored the redox state of DsbA, we observed that cells grown in oleate accumulate DsbA in a completely reduced form during entry into stationary phase (time point T3). However, DsbA resumes to its oxidized form later in the stationary phase (time points T4 and T5) (Figs 1B and 4A). This data suggests that defense mechanisms are induced in oleate-grown cells to deal with the hypo-oxidizing environment of the periplasm. To identify these stress response mechanisms, we checked induction of the five well-characterized ESR pathways in *E. coli* (Bae, Cpx, Psp, Rcs and σ^E^) by assaying β-gal activity of chromosomal *lacZ* fusions to the promoters of their known regulon members (*spy* for Bae, *cpxP* for Cpx, *pspA* for Psp, *rprA* for Rcs and *rpoH*P3 for σ^E^) [2, 3] at different phases of growth (Fig 5A). Since the above reporter fusions were in MG1655 background, we first ensured that this strain cultured in oleate also exhibits inadequate disulfide bond formation. Similar to BW25113, MG1655 cultured in TBK-Ole accumulated a completely reduced form of DsbA at time point T3 (corresponding to entry into stationary phase) (Figs 5A and S2 Fig). Of the five ESR pathways, only Cpx and σ^E^ were upregulated in TBK-Ole grown cells, with Cpx being activated to a greater extent (Figs 5B-F). Importantly, the upregulation of these systems was observed in the stationary phase (Figs 5C and F) and was dependent on oleate utilization (Figs 5G and H). Collectively, these data indicate that Cpx and σ^E^ induction in oleate-utilizing cells might be a combat strategy to maintain redox homeostasis in the envelope.

**Figure 5.**
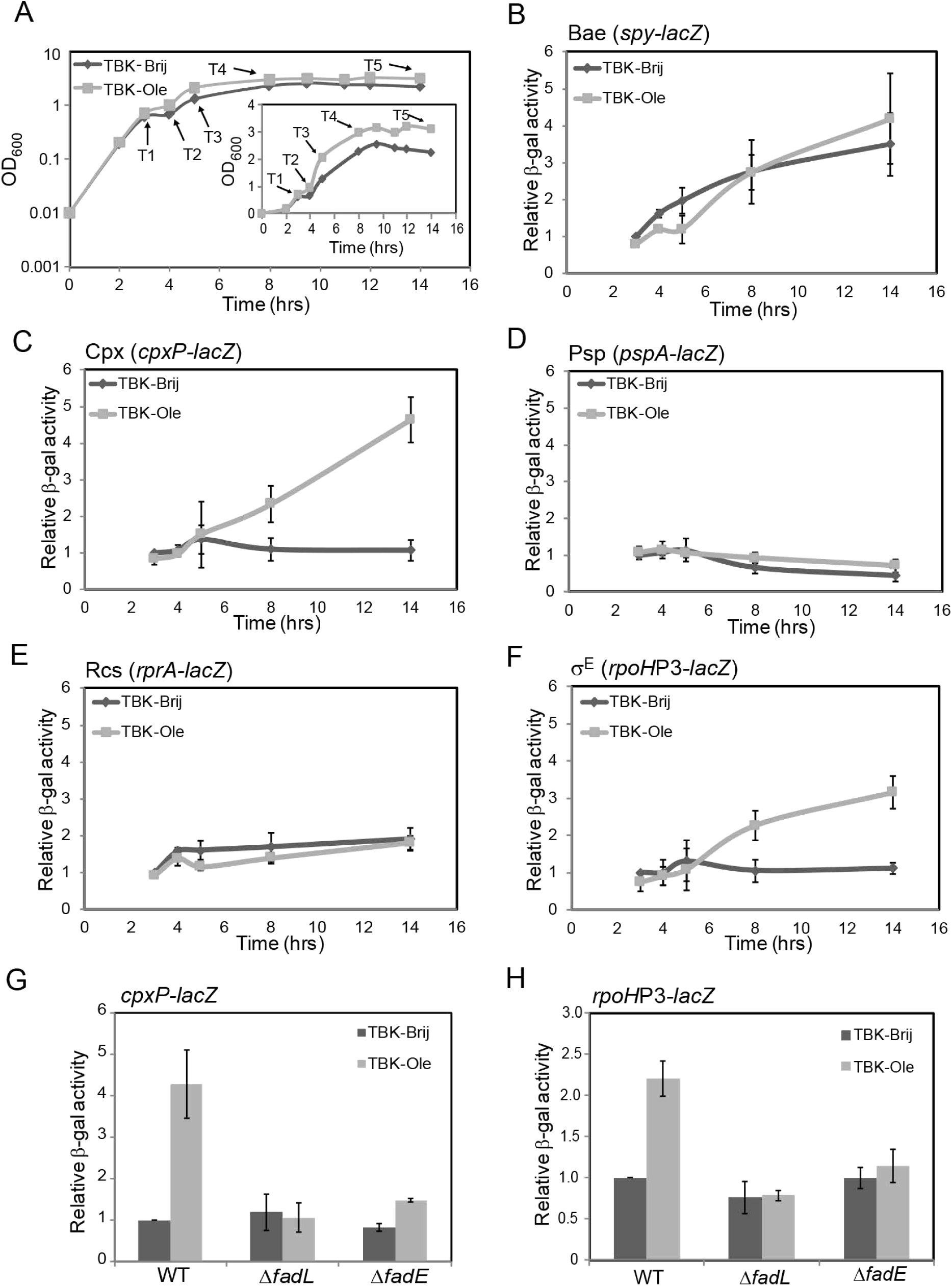
Amongst the various envelope stress response systems in *E. coli*, only Cpx and σ^E^ are induced during LCFA metabolism. *A) Growth curve of WT MG1655 carrying cpxP-lacZ reporter fusion in TBK-Brij and TBK-Ole.* Cells were grown either in TBK-Brij or TBK-Ole. OD_600_ of the cultures was measured, and growth curves were plotted on a semi-logarithmic scale. The experiment was done three times. A representative dataset is shown. T1, T2, T3, T4, and T5 indicate time points where cultures were harvested for β-gal assays. The growth pattern of other reporter strains in MG1655 was similar and thus samples for β-gal assays were collected at the analogous time points. *Inset:* The above growth curves were also plotted on a linear scale. *B-F) Cpx and σ*^*E*^ *responses are induced in stationary phase in E. coli cultured in oleate.* WT MG1655 carrying chromosomal fusion of *lacZ* with the promoters of known regulon members of Bae (*spy-lacZ*) (B), Cpx (*cpxP-lacZ*) (C), Psp (*pspA-lacZ*) (D), Rcs (*rprA-lacZ*) (E), and σ^E^ (*rpoH*P3*-lacZ*) (F) response systems were grown either in TBK-Brij or TBK-Ole. Cultures were harvested at different phases of growth corresponding to the time points indicated in Fig 5A, and β-gal activity was measured. Data were normalized to the β-gal activity of WT in TBK-Brij at time point T1. Data represent the average (± S.D.) of at least three independent experiments. The average β-gal activity of the various reporter strains in TBK-Brij at time point T1 was: 34 (± 4) Miller units for *spy-lacZ*, 41 (± 8) Miller units for *cpxP-lacZ*, 150 (± 75) Miller units for *pspA-lacZ*, 43 (± 2) Miller units for *rprA-lacZ* and 33 (± 15) Miller units for *rpoH*P3*-lacZ. G-H) Activation of Cpx and σ*^*E*^ *responses in E. coli cultured in oleate is dependent on oleate utilization.* Strains carrying *cpxP-lacZ* (G) or *rpoH*P3*-lacZ* (H) reporter fusions were grown either in TBK-Brij or TBK-Ole. Cultures were harvested in the stationary phase corresponding to the time point T5 indicated in Fig 5A, and β-gal activity was measured. Data were normalized to the β-gal activity of WT in TBK-Brij. Data represent the average (± S.D.) of three independent experiments. The average β-gal activity of the WT *cpxP-lacZ* reporter strain in TBK-Brij was 51 (± 9) Miller units and that of the WT *rpoH*P3*-lacZ* was 87 (± 31) Miller units.

### Envelope signals generated by LCFA metabolism activate Cpx and σ^E^

In *E. coli*, the Cpx and σ^E^ systems are induced by a vast array of stressors, including both cytoplasmic and envelope signals. Misfolded proteins, such as the pilin subunits (PapE and PapG) of uropathogenic *E. coli* and the outer-membrane lipoprotein NlpE, constitute the envelope signals for Cpx activation. These signals mediate phosphorylation of an inner-membrane sensor histidine kinase CpxA, which in turn phosphorylates the cytoplasmic response regulator CpxR (Fig 6A, left panel) [32-35]. On the other hand, the low-molecular-weight phosphodonor, acetyl phosphate, a product of the phosphotransacetylase (Pta)-acetate kinase (AckA) pathway constitutes the cytoplasmic signal for Cpx response. In cells grown in glucose, high levels of acetyl phosphate phosphorylate CpxR in a CpxA-independent manner (Fig 6A, right panel) [36]. We observed that in contrast to glucose-grown cells where Cpx induction was majorly in response to the cytoplasmic signal, in oleate-grown cells, Cpx induction was mainly in response to envelope signal. In a Δ*pta*Δ*ackA* mutant where the cytoplasmic Cpx signal is eliminated, Cpx induction reduced to ∼30% of WT in glucose; however, it decreased to only ∼70% of WT in oleate. On the other hand, in a Δ*cpxA* mutant where Cpx signaling from the envelope is eliminated, Cpx induction reduced to ∼30% of WT in oleate; however, it decreased to only ∼70% of WT in cells grown in glucose (Figs 6B and C). The Δ*pta*Δ*ackA* strain exhibited lower Cpx induction compared to WT in TBK-Brij likely due to the elimination of signaling from acetyl-phosphate generated during growth in this medium (*Inset* Fig 6B). On the other hand, the Δ*cpxA* strain showed higher Cpx induction in basal medium, likely due to the high steady-state levels of phosphorylated CpxR (phosphorylated by acetyl phosphate) resulting from the loss of phosphatase activity of CpxA (*Inset* Fig 6C) [36, 37].

**Figure 6.**
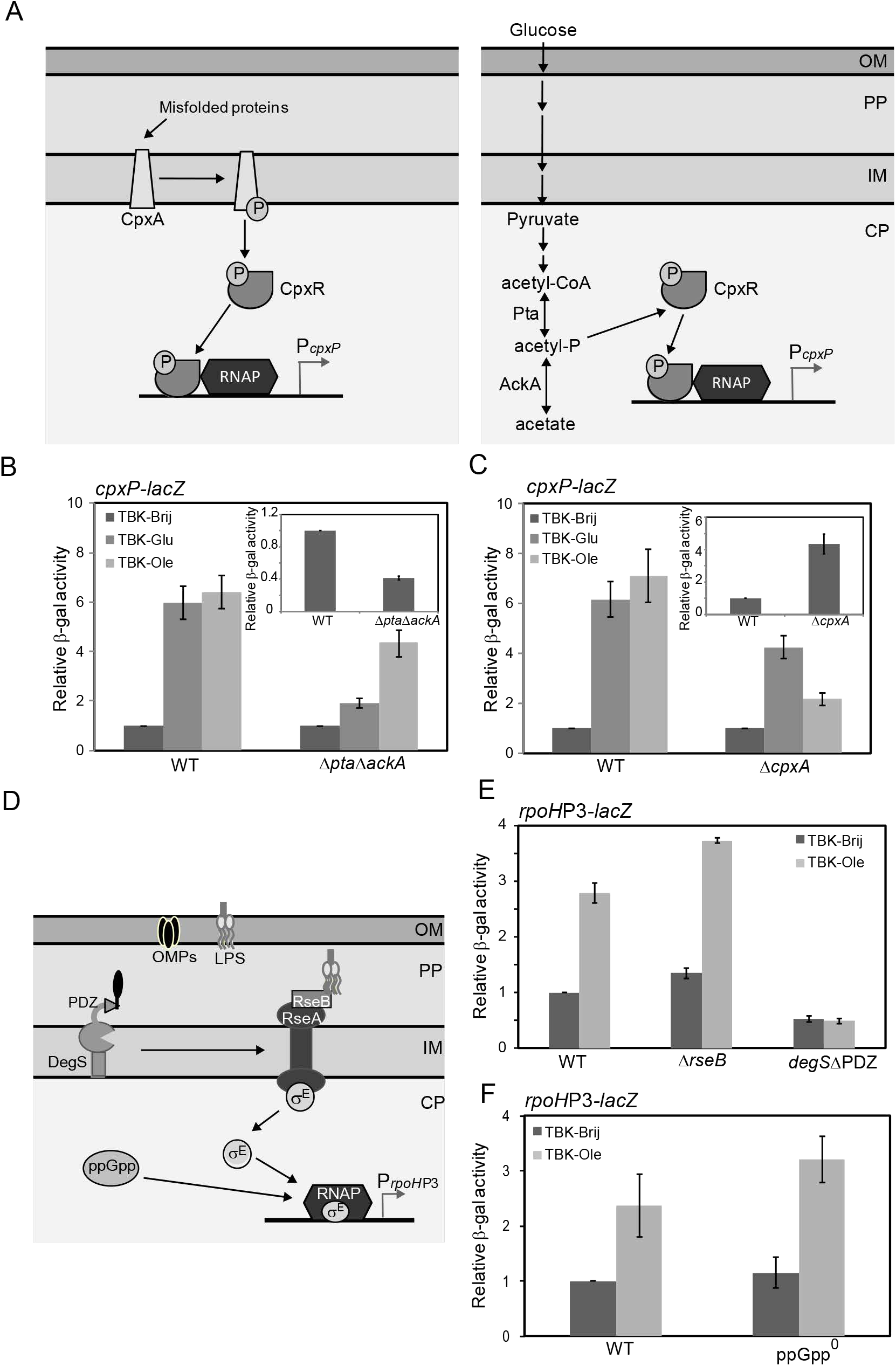
Cpx and σ^E^ pathways are activated in response to envelope signals generated by LCFA metabolism. *A) Organization of the Cpx pathway.* The CpxAR two-component system comprises of an inner-membrane sensor histidine kinase, CpxA and a cytoplasmic response regulator, CpxR. <u>Left panel:</u> In the presence of envelope signals, such as misfolded proteins, CpxA acts as a kinase and phosphorylates CpxR. <u>Right panel:</u> Acetyl-phosphate, an intermediate of the Pta-AckA pathway, acts as the cytoplasmic signal for Cpx activation in cells grown in glucose, by directly transferring its phosphoryl group to CpxR. Phosphorylated CpxR directs the transcription of genes involved in mitigating stress. Abbreviations: CP, cytoplasm; IM, inner membrane; PP, periplasm; OM, outer membrane; acetyl-P, acetyl-phosphate; RNAP, RNA Polymerase. *B and C) Induction of the Cpx pathway in oleate-grown cells is majorly in response to envelope signal(s).* Strains carrying *cpxP-lacZ* reporter fusion were grown in TBK-Brij, TBK-Brij supplemented with glucose (TBK-Glu) or TBK-Ole. Cultures were harvested in the stationary phase, and β-gal activity was measured. Data obtained for TBK-Glu and TBK-Ole for each strain were normalized to the β-gal activity of that strain in TBK-Brij and represent the average (± S.D.) of three independent experiments. *Insets:* The data from the main Figs 6B and C are plotted as β-gal activity of deletion strain in TBK-Brij normalized to the β-gal activity of WT in the same medium. The average β-gal activity of the WT *cpxP-lacZ* reporter strain in TBK-Brij was 37 (± 5) Miller units (B) and 26 (± 3) Miller units (C). *D) Organization of the σ*^*E*^ *pathway.* Under unstressed conditions, σ^E^ remains bound to RseA and is thus less available to bind RNA polymerase. Unfolded OMPs activate DegS by binding to its PDZ domain. LPS displaces RseB from RseA enabling activated DegS to cleave RseA in its periplasmic domain. RseA is finally degraded by additional proteases to release σ^E^, which binds RNA polymerase and initiates transcription of its regulon members. The cytoplasmic alarmone ppGpp upregulates σ^E^-dependent transcription. Abbreviations: CP, cytoplasm; IM, inner membrane; PP, periplasm; OM, outer membrane; LPS, lipopolysaccharide; OMPs, outer membrane proteins; RNAP, RNA Polymerase. *E and F) Induction of the σ*^*E*^ *pathway in oleate-grown cells is in response to envelope signal(s).* Strains carrying *rpoH*P3*-lacZ* were grown either in TBK-Brij or TBK-Ole. Cultures were harvested in the stationary phase and β-gal activity was measured. Data were normalized to the β-gal activity of WT in TBK-Brij and represent the average (± S.D.) of three independent experiments. The average β-gal activity of the WT *rpoH*P3*-lacZ* in TBK-Brij was 29 (± 3) Miller units (E) and 71 (± 9) Miller units (F).

The two major components of the outer membrane, OMPs and LPS, constitute the envelope signals for σ^E^ induction. Unfolded OMPs activate the inner membrane-anchored periplasmic protease DegS by binding to its PDZ domain. Activated DegS initiates cleavage of the anti-σ^E^ factor RseA, which normally sequesters σ^E^ at the inner membrane and prevents it from binding RNA polymerase [38]. LPS activates σ^E^ by displacing the periplasmic protein, RseB away from RseA, enabling OMP-activated DegS to cleave RseA [39]. The σ^E^ response is also activated independent of the envelope signals. Under nutritional stress, a cytoplasmic alarmone factor, guanosine 3’,5’-bispyrophosphate (ppGpp), upregulates σ^E^-dependent transcription [40] (Fig 6D). We observed that in oleate-grown cells, σ^E^ induction was completely abolished in a *degS*ΔPDZ strain; however, σ^E^ activation was not affected either in a Δ*rseB* strain or a ppGpp^0^ strain that lacks enzymes (RelA and SpoT) involved in ppGpp synthesis (Figs 6E and F). The requirement of the DegS PDZ domain strongly suggests that the signal for σ^E^ induction during oleate metabolism is an unfolded OMP. Taken together, our data indicate that both Cpx and σ^E^ pathways are activated in response to envelope signals generated by LCFA metabolism.

### The hypo-oxidizing environment of the envelope is one of the reasons for Cpx induction during LCFA metabolism

We showed above that LCFA metabolism results in insufficient disulfide bond formation and generates envelope signals for the activation of Cpx and σ^E^ pathways. We next investigated whether these signals are generated because of the hypo-oxidizing environment of the envelope. To test this, we measured Cpx and σ^E^ induction in oleate-grown cells upon exogenous supplementation of ubiquinone, which would make the environment of the periplasm oxidizing by taking up electrons from the disulfide bond-forming machinery. Whereas Cpx induction decreased by ∼40% in TBK-Ole grown cells, there was no effect on σ^E^ induction (Fig 7A). This data indicates that during LCFA metabolism, the hypo-oxidizing environment of the envelope generates a stressor for Cpx activation, whereas the signal for σ^E^ induction is likely redox-independent. As a separate test that the Cpx pathway is activated in response to the altered redox state of the envelope, we used the Δ*cydD* strain, which is defective in exporting reduced glutathione and cysteine to the periplasm and thus results in a hyper-oxidizing envelope [41, 42]. Cpx response was downregulated ∼2-fold in the Δ*cydD* strain in both basal and oleate supplemented media; however, there was no effect on σ^E^ induction (Fig 7B). Finally, we checked the induction of Cpx and σ^E^ upon supplementation of TBK-Brij medium with DTT, which would outcompete the oxidation of periplasmic proteins by DsbA resulting in accumulation of reduced proteins in the periplasm [31]. Whereas there was ∼6 fold induction of the Cpx response at 3 mM DTT, σ^E^ was induced only ∼2-fold (S3 Fig). Collectively, our data emphasize that Cpx is the major pathway that gets activated in response to the altered redox state of the envelope.

**Figure 7.**
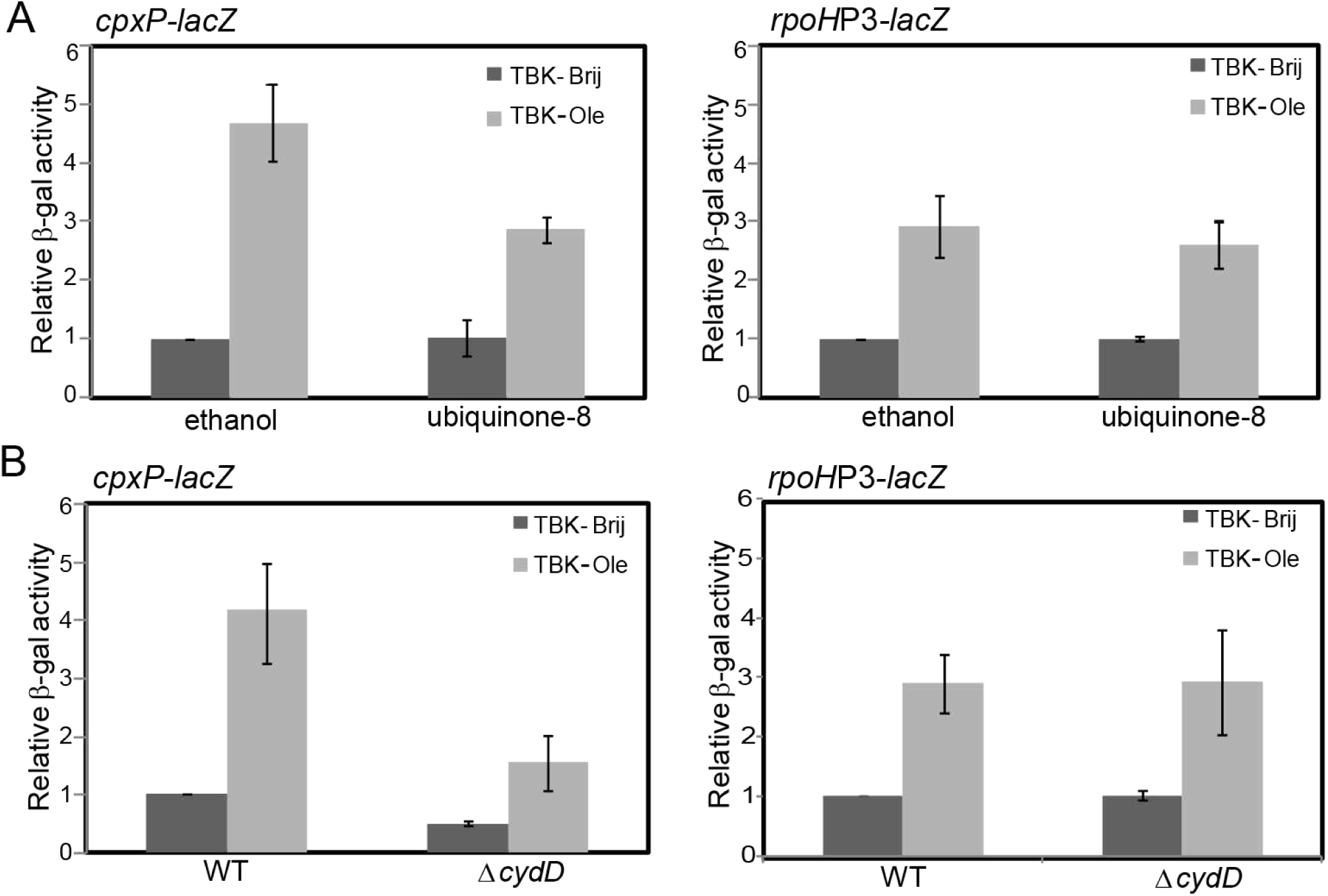
During LCFA metabolism Cpx is induced partly in response to the hypo-oxidizing environment of the envelope. *A) Supplementation of ubiquinone-8 decreases Cpx induction in oleate-utilizing cells.* WT carrying either *cpxP-lacZ* (left panel) or *rpoH*P3*-lacZ* (right panel) reporter fusion was grown either in TBK-Brij or TBK-Ole. The media contained either 20 μM ubiquinone-8 or 0.1% ethanol. Cultures were harvested in the stationary phase, and β-gal activity was measured. Data were normalized to the β-gal activity of WT in TBK-Brij supplemented with 0.1% ethanol and represent the average (± S.D.) of three independent experiments. The average β-gal activity of the *cpxP-lacZ* reporter strain in TBK-Brij supplemented with 0.1% ethanol was 30 (± 13) Miller units and that of the *rpoH*P3*-lacZ* was 57 (± 17) Miller units. *B) Cpx response is downregulated in a* Δ*cydD strain.* Strains carrying either *cpxP-lacZ* (left panel) or *rpoH*P3*-lacZ* (right panel) reporter fusion were grown either in TBK-Brij or TBK-Ole. Cultures were harvested in the stationary phase, and β-gal activity was measured. Data were normalized to the β-gal activity of WT in TBK-Brij and represent the average (± S.D.) of three independent experiments. The average β-gal activity of the WT *cpxP-lacZ* reporter strain in TBK-Brij was 53 (± 12) Miller units and that of the WT *rpoH*P3*-lacZ* was 51 (± 16) Miller units.

### Cpx and σ^E^ pathways maintain proper disulfide bond formation in the envelope

Our above results suggest that the activation of both Cpx and σ^E^ responses during LCFA metabolism occurs due to stress signals from the envelope; however, only Cpx is activated in response to redox imbalance. Next, we investigated whether the activation of these pathways helps in managing disulfide bond formation. For this, we determined the effect of preventing the induction of Cpx regulon members (by using Δ*cpxR* strain) and of downregulating σ^E^ (by using the *degS*ΔPDZ strain where σ^E^ activity is downregulated ∼2-fold; Fig 6E and [39]) on AP activity in TBK-Brij and TBK-Ole media. Whereas there was no effect of downregulation of σ^E^ on AP activity in either of the media conditions, we observed ∼40% decrease in AP activity in a Δ*cpxR* strain in the TBK-Brij medium, an effect similar to that observed in WT cultured in TBK-Ole (Fig 8). This result suggests that preventing Cpx activation hampers disulfide bond formation even in a basal medium, which could be due to the downregulation of DsbA, a Cpx regulon member directly involved in disulfide bond formation [43]. We did not observe a further decrease in AP activity in a Δ*cpxR* strain grown in the presence of oleate. This is likely because the disulfide bond-forming machinery is already compromised in oleate-utilizing cells, and thus a decrease in DsbA levels due to *cpxR* deletion would not decrease AP activity further.

**Figure 8.**
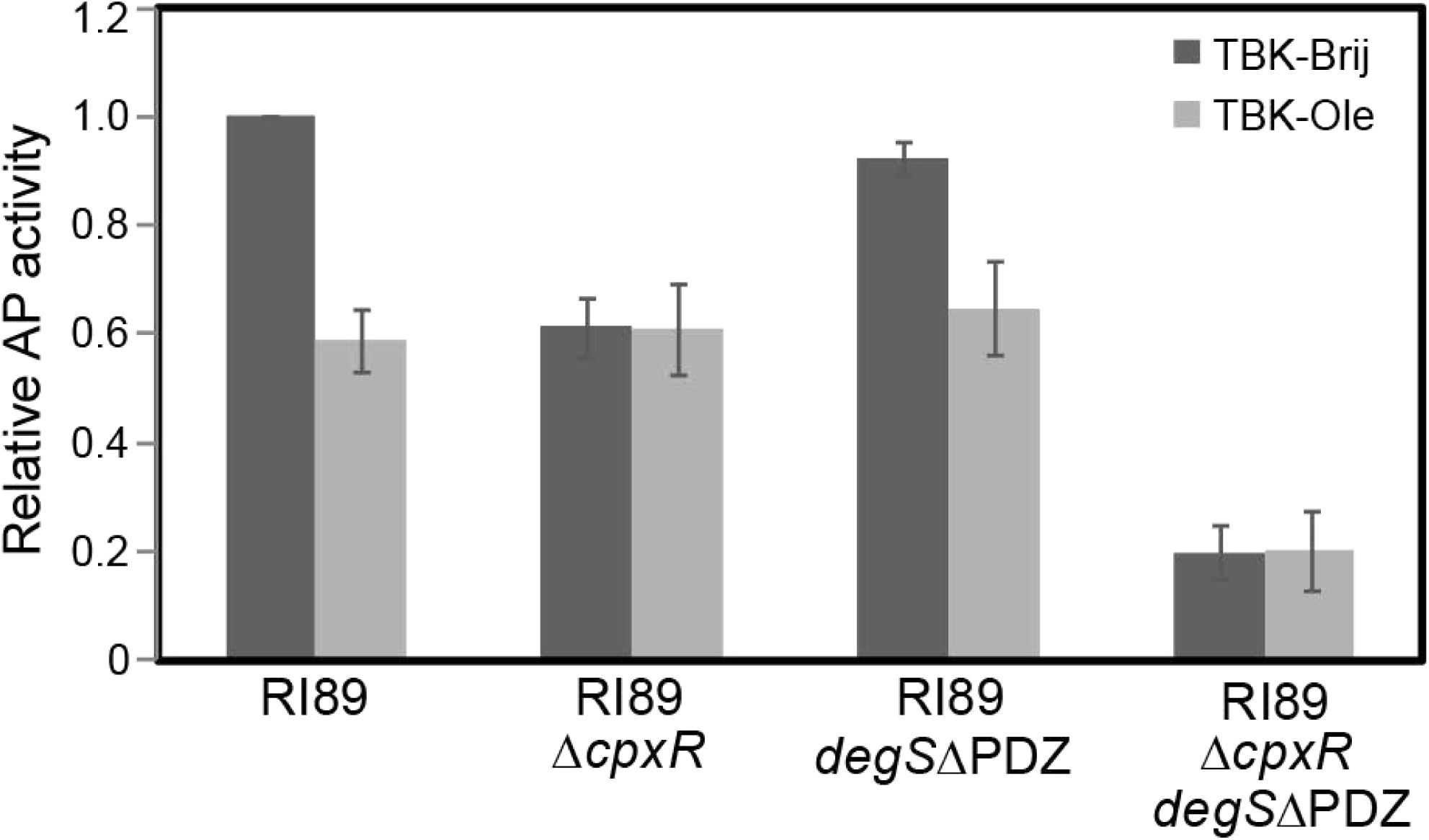
Cpx and σ^E^ pathways maintain proper disulfide bond formation. Strains were grown either in TBK-Brij or TBK-Ole. Cultures were harvested in the exponential phase, and AP activity was measured. Data were normalized to the AP activity of RI89 in TBK-Brij and represent average (± S.D.) of three independent experiments. The average AP activity of RI89 in TBK-Brij was 1346 (± 233) units.

σ^E^ regulates several periplasmic chaperones and proteases [44]; therefore, we investigated if the σ^E^ response prevents overburdening of the disulfide bond-forming machinery in a Δ*cpxR* strain by repairing or clearing out unfolded/ misfolded proteins that accumulate in this strain due to compromised disulfide bond formation. To test this, we measured AP activity of the Δ*cpxR degS*ΔPDZ strain in TBK-Brij and TBK-Ole. Consistent with our proposal, we find that the downregulation of σ^E^ activity in a Δ*cpxR* background reduces AP activity to ∼20% of the WT cultured in TBK-Brij (Fig 8). Collectively, these results suggest that whereas the Cpx pathway plays a major role in maintaining disulfide bond formation, the σ^E^ response plays an ancillary role in this process by managing damaged proteins.

## Discussion

Here we investigated the interconnection between LCFA metabolism, envelope redox status, and ESR pathways. We demonstrated that during LCFA metabolism, increased electron flow in the ETC, renders ubiquinone limiting for disulfide bond formation, thereby compromising redox balance in the envelope. Signals generated during growth in LCFAs activate Cpx and σ^E^ ESR systems, which maintain redox homeostasis. Importantly, our work suggests that of the two ESR systems, Cpx plays a primary role in this process: *i)* amongst Cpx and σ^E^, Cpx is induced to a greater extent by LCFAs, *ii)* only Cpx pathway senses redox-dependent signal, and *iii)* σ^E^ downregulation hampers disulfide bond formation only in the absence of Cpx response.

### Ubiquinone is limiting during LCFA metabolism

Ubiquinone has been considered non-limiting for its electron transfer function based on an earlier observation that during aerobic growth of *E. coli* (in minimal medium supplemented with glucose), it is present in ∼15 to 20-fold excess over flavins and cytochromes [45]. However, our previous work suggested that ubiquinone is limiting during aerobic metabolism of LCFAs. Briefly, we showed that LCFA degradation generates high levels of reactive oxygen species (ROS) and ubiquinone counteracts this oxidative stress. We proposed that high NADH/NAD^+^ and FADH_2_/FAD ratios during LCFA catabolism increase electron flow in the ETC, thereby increasing ROS formation, and ubiquinone decreases ROS by rapidly transferring electrons from the site of ROS formation to the ETC. Despite the endogenous accumulation of ubiquinone (∼2-fold), its exogenous supplementation was still required to decrease ROS in LCFA-utilizing cells. These data strongly suggested that ubiquinone is limiting under metabolic conditions that feed a large number of electron donors in the ETC [23]. Here we show that electron flow in the ETC is indeed high during LCFA metabolism (Fig 1), and in addition to being insufficient for transfer of electrons derived from metabolism, ubiquinone is also limiting for electron transfer from the disulfide bond-forming machinery. The latter is evident from several characteristics of inadequate disulfide bond formation exhibited by LCFA-utilizing cells, which are prevented upon ubiquinone supplementation (Figs 2-4 and S1 Fig). Although compromised disulfide bond formation has been observed earlier under conditions that result in either defective or non-operational ETC [16, 17], aerobic metabolism of LCFAs represents the first instance, where in an otherwise operational ETC, disulfide bond formation is hampered due to an increased electron flow from carbon metabolism.

We observed accumulation of the reduced form of DsbA when LCFA-grown cells entered the stationary phase, which restored to the oxidized state in mid- and late-stationary phases of growth (Fig 4A). A previous study showed that during aerobic growth of *E. coli* (in minimal medium supplemented with glucose) an increase in cell density increases the ubiquinol/ubiquinone ratio due to a decrease in the availability of dissolved oxygen, and this ratio decreases later in stationary phase due to lowered metabolism [46]. Because LCFA metabolism produces a large number of electron donors, this would lead to a much larger increase in ubiquinol/ubiquinone ratio with an increase in cell density, thereby making ubiquinone less available for the oxidation of the disulfide bond-forming machinery. The restoration of the oxidized form of DsbA in the stationary phase would be an outcome of a decrease in the ubiquinol/ubiquinone ratio due to the downregulation of LCFA metabolism. We suggest that Cpx activation in LCFA-utilizing cells helps restore redox homeostasis partly by downregulating metabolism (see below).

### Possible mechanisms by which Cpx and σ^E^ sense envelope stress during LCFA metabolism

Several lipoproteins, misfolded pilin subunits of uropathogenic *E. coli*, and inner membrane respiratory complexes are known inducers of the Cpx pathway [33, 35, 47, 48]. Although the details of Cpx activation by these envelope components remains largely elusive, there are two suggested mechanisms. One mechanism involves direct interaction of the molecular signal with the sensor kinase, CpxA. An outer membrane lipoprotein, NlpE, accumulates at the inner membrane due to defects in lipoprotein trafficking and physically interacts with CpxA. Since NlpE is also a DsbA substrate, its mutant lacking the C-terminal disulfide bond turns on Cpx and Cpx activation in Δ*dsbA* is NlpE-dependent, this lipoprotein has been proposed to sense redox imbalance in the envelope [49, 50]. Another mechanism of Cpx activation involves titration of CpxP, a periplasmic inhibitor of CpxA. Misfolded pilin subunits sequester CpxP and deliver it to the periplasmic protease DegP. In this process, DegP degrades both CpxP and the pilin subunits [35]. However, CpxP displacement itself is not a sensing mechanism since pilin subunits activate Cpx even in its absence. CpxP is therefore proposed to fine-tune Cpx response by preventing inappropriate activation of CpxA and enabling rapid Cpx downregulation once the envelope stress is mitigated [51].

The utilization of LCFAs generates envelope signals for Cpx activation, which are likely both redox-dependent and redox-independent. The presence of two types of signals is supported by the observation that upon providing ubiquinone to oleate-grown cells, whereas DsbA is present in its oxidized form and there is no decrease in AP activity, Cpx response is only partially downregulated (Figs 2D, 4B, and 7A). Because NlpE is a well-recognized molecular signal for Cpx and can also sense redox imbalance, we tested if it is the molecular cue during LCFA metabolism. However, we find that Cpx is fully induced in a Δ*nlpE* strain grown in oleate (S4 Fig). Because DegP is also a DsbA substrate and functions as a protease in the thiol state [30], it is plausible that during LCFA metabolism, DegP accumulates in a reduced form and degrades CpxP. However, since CpxP titration only fine-tunes the Cpx response, the additional redox-independent signal likely interacts directly with CpxA, resulting in robust activation of Cpx. Alternatively, both redox-dependent and redox-independent signals might induce Cpx exclusively via CpxA, without involving CpxP. It will be worth testing whether respiratory complexes or lipoproteins other than NlpE constitute signal(s) for Cpx activation during growth in LCFAs.

The σ^E^ pathway senses perturbations in the biogenesis of the outer membrane components, OMPs and LPS, via the DegS PDZ domain and RseB, respectively [5, 38, 39]. In a Δ*dsbA* strain, σ^E^ activation does not require the DegS PDZ domain and is suggested to be via RseB inhibition. An outer membrane component of the LPS transport machinery, LptD, is a DsbA substrate. Therefore, it is suggested that a decrease in the level of properly disulfide-bonded LptD in a Δ*dsbA* strain results in LPS accumulation, which then inhibits RseB to activate σ^E^ [39]. Although LCFA metabolism also causes redox imbalance, σ^E^ induction is likely in response to redox-independent envelope signal (Fig 7A). Further, the complete abolishment of σ^E^ induction in a *degS*ΔPDZ strain, but not in a Δ*rseB* strain, strongly indicates that an unfolded OMP, and not LPS, is the molecular signal for σ^E^ activation (Fig 6E). The probable candidate is FadL, the outer membrane LCFA transporter, which is upregulated during growth in LCFAs [23], and activates σ^E^ when overexpressed [52]. Taken together, multiple cues induce Cpx and σ^E^ pathways, which differ in their mode of activation. We suggest that even for conditions that ultimately impact the redox status of the envelope, i.e., deletion of *dsbA* and LCFA metabolism, the molecular signals which activate these stress responses are distinct.

### Probable feedback exerted by Cpx and σ^E^ to maintain envelope redox homeostasis during growth in LCFAs

The Cpx and σ^E^ responses mediate adaptation to damaged proteins by upregulating envelope-localized protein folding and degrading factors [4, 44, 53]. Because insufficient disulfide bond formation in LCFA-grown cells will result in the accumulation of unfolded/ misfolded proteins, the induction of Cpx and σ^E^ might restore envelope homeostasis by repairing/ clearing out damaged proteins. DsbA, a Cpx regulon member, maintains redox homeostasis by forming disulfide bonds in thiol proteins [11, 43]. However, during growth in LCFAs, DsbA upregulation might exacerbate the situation because newly synthesized DsbA would itself require disulfide bond formation for its activity. A more effective mechanism for restoring redox homeostasis would be to downregulate components of the metabolic pathway that feed electrons into the ETC and upregulate components that increase the oxidizing power of the ETC. In fact, a combination of the following observations suggests that Cpx pathway can maintain redox homeostasis by modulating LCFA metabolism: *i)* Cpx strongly downregulates NADH dehydrogenase I and succinate dehydrogenase, which transfer electrons into the ETC [47, 53], and *ii)* in the Δ*cydD* strain, which has a hyper-oxidizing envelope, Cpx is downregulated (Fig 7B; this study), whereas *fad* genes involved in β-oxidation are upregulated [54]. Future studies are needed to investigate the regulation exerted by Cpx on the LCFA metabolic pathway. These investigations will unravel that besides repairing envelope by dealing with damaged proteins, Cpx also prevents envelope redox stress by modulating electron flow from LCFA metabolism, providing further support to our proposal that Cpx plays a primary role in maintaining redox homeostasis. Figure 9 summarizes the findings from the current work, and presents model for Cpx and σ^E^ activation during LCFA metabolism and the feedback provided by these systems to maintain envelope redox homeostasis.

**Figure 9.**
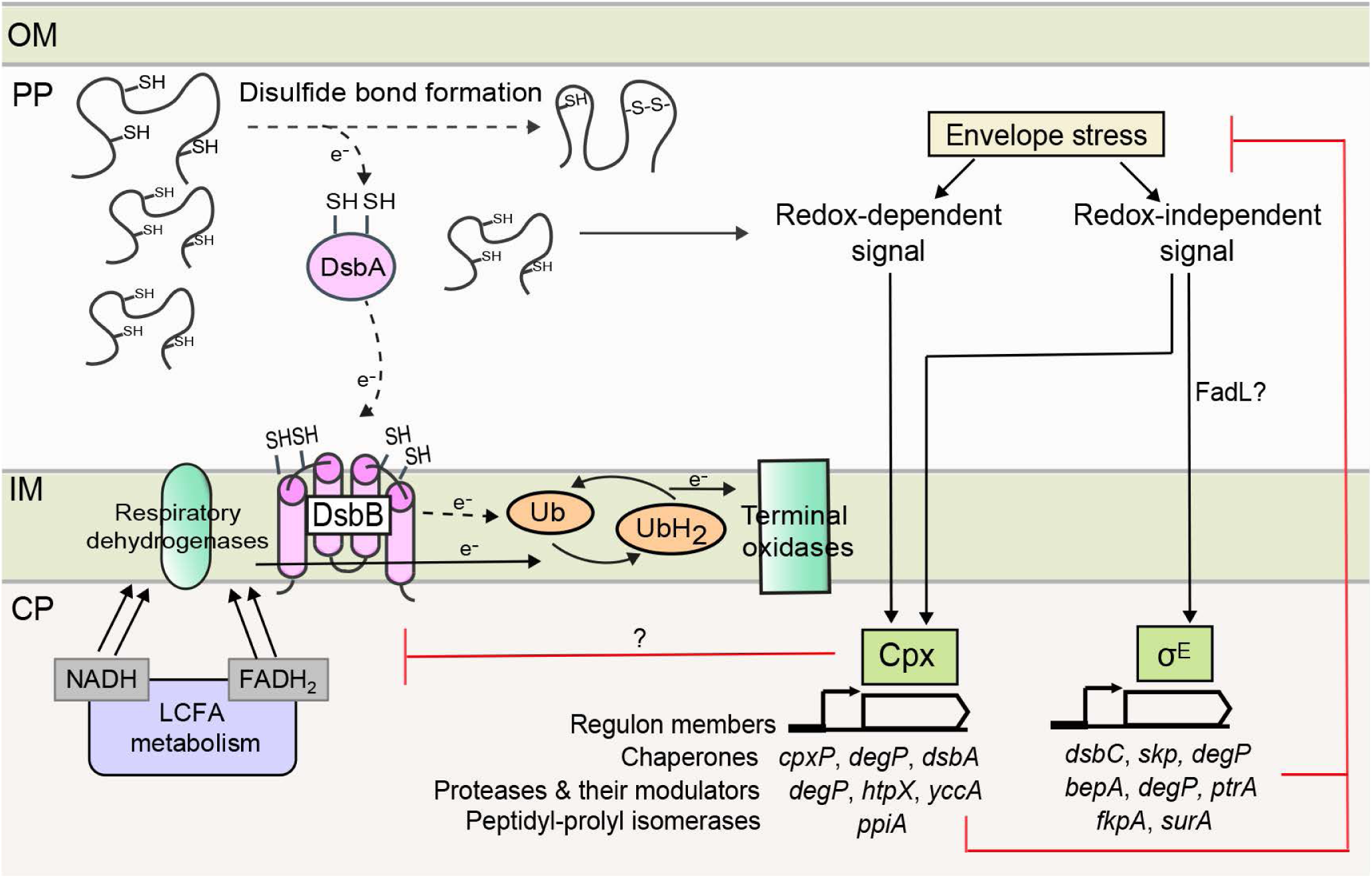
Model depicting the interconnection between LCFA metabolism, envelope redox status, and ESR pathways. LCFA degradation produces a large number of NADH and FADH_2_, which are oxidized by respiratory dehydrogenases in the ETC. The increased flow of electrons in the ETC makes ubiquinone insufficient for taking up electrons from DsbA-DsbB, resulting in the accumulation of unfolded/ misfolded proteins. Cpx and σ^E^ ESR systems are activated, which restore envelope homeostasis. Cpx is partly induced by a redox-dependent signal, which is likely a DsbA substrate. Some redox-independent signal also activates Cpx. σ^E^ is activated in response to the accumulation of an unfolded OMP, likely FadL, and this constitutes its redox-independent signal. Cpx and σ^E^ responses decrease envelope stress by upregulating chaperones, proteases and their modulators, and peptidyl-prolyl isomerases, which repair damaged proteins. Besides, Cpx might decrease the load on ETC by downregulating LCFA metabolism. Arrows with *e*^-^ labeled on the line denote the direction of electron flow. The dotted arrows indicate decreased electron flow at these steps. Abbreviations: CP, cytoplasm; IM, inner membrane; PP, periplasm; OM, outer membrane; Ub, ubiquinone; UbH_2_, ubiquinol.

Several gram-negative bacterial pathogens such as *Pseudomonas aeruginosa, Salmonella typhimurium*, and *Vibrio cholerae* use LCFAs acquired from host tissues as their source of energy [19-21]. The present study provides a rationale for examining whether LCFA metabolism induces redox stress in these pathogens and whether activation of ESR pathways equivalent of Cpx and σ^E^ represents their combat strategy. In addition to LCFA metabolism, disulfide bond formation and ESR systems have also been associated with bacterial pathogenesis [19-21, 55-62]. Therefore, it will be important to determine whether LCFA metabolism impacts pathogenesis by influencing the redox status of the envelope and activation of ESR pathways.

## Materials and methods

### Strains, plasmids, and primers

Strains and plasmids are listed in S1 Table. Primers used in this study are listed in S2 Table. For deletion strains obtained from the Keio collection [63], their fresh transductants were made by P1 transduction and verified by colony PCR to rule out genetic errors. The derivatives of *degS*ΔPDZ strain were unstable as glycerol stocks and thus were always freshly made before use.

To determine the induction of *fadE*, chromosomal single-copy transcriptional reporter fusions were made using the pAH125 integration vector, as described previously [64]. Briefly, 980bp DNA fragment upstream of the start codon of *fadE* gene was amplified from the genomic DNA of BW25113 and cloned upstream of *lacZ* at the *Kpn*I and *Eco*RI sites of pAH125. The resulting construct was integrated at the *attλ* site of BW25113 using integrase expressed from the pINT-ts helper plasmid. Single-copy integrants were verified by PCR. Reporter integrations were P1 transduced into a clean BW25113 strain.

The transcriptional reporter fusion for the Psp ESR pathway, RC15106, was constructed as follows. P1 lysate from AM1247 was used to infect MG1655Δ*lac* strain. Transductants were screened for Lac+ phenotype on LB agar containing 10 mM citrate and 0.1 mg/ml X-gal. Colonies were re-streaked to obtain pure blue colonies. Blue colonies were further screened for the presence of *nadA* based on tetracycline sensitivity.

The ppGpp^0^ strain was always freshly made before use. Briefly, RC15105 was constructed by transducing *relA::kan* from the Keio collection into CAG45114. The *spoT::cam* allele from CAG55907 was then transduced into RC15105. The ppGpp^0^ phenotype was confirmed by the inability of the strain to grow on M9 minimal medium lacking amino acids [40]. Construction of ppGpp^0^ strain and experiments involving this strain were performed at 30°C.

### Media composition and growth conditions

Media had the following composition: lysogeny broth (LB) was 5 g/liter Bacto yeast extract, 10 g/liter Bacto tryptone, and 5 g/liter NaCl; tryptone broth K (TBK) was 10 g/liter Bacto tryptone, and 5 g/liter KCl [36]; M9 minimal medium was 5.3 g/liter Na_2_HPO_4_, 3 g/liter KH_2_PO_4_, 0.5 g/liter NaCl, 1 g/liter NH_4_Cl, 0.12 g/liter MgSO_4_, 2 mg/liter biotin, 2 mg/liter nicotinamide, 0.2 mg/liter riboflavin, and 2 mg/liter thiamine. TBK media was buffered at pH 7.0 using 100 mM potassium phosphate buffer. Where required, media were supplemented with either glucose or sodium salt of acetate or oleate, at a final concentration of 5 mM. Stock of oleate (50 mM) was prepared in 5.0% Brij-58 [23]. Media were solidified using 1.5% (w/v) Difco agar. When required, ampicillin (100 μg/ml), chloramphenicol (20 μg/ml), kanamycin (10 or 30 μg/ml) and tetracycline (25 μg/ml) were used.

Primary cultures were grown in 3 ml LB. Secondary cultures were set-up either in TBK or in M9 minimal medium containing the desired carbon source and grown for defined time periods. When required, CdCl_2_, DTT, 1-thioglycerol (at desired concentrations) and ubiquinone (20 μM) were added to the medium. Except for ppGpp^0^, all cultures were incubated at 37°C.

### Growth curves in shake flasks and 96-well plates

Growth curves in TBK medium were performed in shake flasks. Primary cultures were pelleted, washed, and re-suspended in TBK medium. Cells were re-inoculated in 15 ml medium in 125 ml flasks to an initial OD_600_ of ∼0.01. Cultures were grown at 37°C, and OD_600_ was measured at definite time intervals.

Growth curves in M9 minimal medium were performed in 96-well plates, as described previously [23]. Briefly, cells washed and re-suspended in M9 minimal medium were re-inoculated using a robotic liquid handling system (Tecan) in 200 μl of the minimal medium containing the desired carbon source to an initial OD_450_ of ∼0.03. Plates were incubated at 37°C in a shaker, and OD_450_ of the cultures was measured at regular time intervals (Tecan Infinite M200 monochromator). The microplate reader and incubator shaker were integrated with the liquid handling system where the transfer of plates between the shaker and reader was automated.

### NADH and NAD^+^ quantitation

The extraction of NAD^+^ and NADH was carried out following the procedure described in [65], with slight modifications. Briefly, 6 ml cultures were pelleted, washed three times with 1 ml cold 1X PBS (8 g/liter NaCl, 0.2 g/liter KCl, 1.44 g/liter Na_2_HPO_4_ and 0.24 g/liter KH_2_PO_4_; pH 7.4), and normalized to OD_450_ ∼1. Immediately, 1 ml of each sample was taken in two microcentrifuge tubes (MCTs), pelleted and re-suspended in 300 μl of 0.2 M NaOH (for NADH) or 0.2 M HCl (for NAD^+^). The samples were incubated at 50°C in a water bath for 10 min and then immediately transferred to ice for 5 min. Following this, 300 μl of 0.1 M HCl (for NADH) or 0.1 M NaOH (for NAD^+^) was added drop-wise to the samples with vortexing. Samples were centrifuged at 18,400 X g for 5 min at 4°C to remove cell debris. The supernatant was transferred to a fresh MCT and kept in ice. Samples were deproteinized using a 10 kDa cut-off filter by centrifugation at 6,900 X g for 15 min at 4°C and kept in ice. The amount of NADH and NAD^+^ in samples was quantitated using the NAD/NADH quantitation kit (Sigma). Briefly, 150 μl reaction mix was prepared in 96-well plates (transparent with clear bottom). Each reaction mix contained 50 μl of the extracted sample, 98 μl cycling buffer, and 2 μl cycling enzyme mix. The reaction mixture was kept at room temperature for 10 min followed by the addition of 10 μl NADH developer in the dark. Samples were incubated further for one hour. Absorbance was measured at 450 nm (Thermo Scientific Multiskan Go). A standard curve was generated using the NADH standard provided in the kit, and the amount of NADH and NAD^+^ in the samples was determined.

### Enzyme activity assays

#### Preparation of cell extract

Cultures were washed at least three times with assay buffer. ∼10^10^ cells were re-suspended and sonicated. Samples were centrifuged at 18,400 X g for 40 min at 4°C. The supernatant was collected and kept in ice. Protein in the cell extracts was quantified using Bradford assay.

#### NADH dehydrogenase assay

The protocol adapted from [66] was slightly modified. 1 ml reaction mixture contained 50 mM Tris-Cl (pH 8.0), 250 μM menadione, and 1 μg protein. The reaction was initiated by adding 250 μM NADH. Enzyme activity was calculated from the decrease in absorbance of NADH (extinction coefficient: 6.22 mM^-1^ cm^-1^) at 340 nm over a period of 5 min. The activity was expressed as nmoles of NADH oxidized per min per mg protein. The reaction mixture without NADH was taken as blank.

#### Succinate dehydrogenase assay

The procedure from [67] was followed with few modifications. 1 ml reaction mixture contained 0.15 M NaPO_4_ (pH 7.0), 0.1 M sodium succinate (pH 7.5), 0.02 M sodium azide, 0.1 mM ubiquinone-2 (Sigma; 10 mM stock was prepared in ethanol) and 50 μg protein. The reaction was followed by adding 0.05 mM DCPIP (2,6-Dichlorophenolindophenol). Enzyme activity was calculated from the decrease in absorbance of DCPIP (extinction coefficient: 22 mM^-1^ cm^-1^) at 600 nm over a period of 15 min. The activity was expressed as nmoles of DCPIP reduced per min per mg protein. The reaction mixture without DCPIP was taken as blank.

### β-galactosidase assay

β-gal assays were performed as mentioned in [23]. Briefly, cells were pelleted, washed four times with Z-buffer, and re-suspended in the same buffer to OD_450_ ∼0.5. Promoter activity was measured by monitoring β-gal expression from chromosomal transcriptional reporter fusions, as described [68].

### Alkaline phosphatase assay

AP activity was determined as described in [69], with slight modifications. Freshly prepared iodoacetamide (1 mM) was added directly to the cultures. Samples were kept on ice for 15 min. Cells were pelleted and washed three times with a buffer comprised of 10 mM Tris-Cl (pH 8.0), 10 mM MgSO_4_, and 1 mM iodoacetamide. The cells were resuspended in a buffer containing 1 M Tris-Cl (pH 8.0), 0.1 mM ZnCl_2_, 0.002% SDS and 20 μl chloroform. The assay was performed at 37°C. The reaction was initiated by the addition of 200 μl of 4 mg/ml *p*-nitrophenyl phosphate and was stopped by the addition of 120 μl of a buffer containing 165 mM K_2_HPO_4_ and 80 mM EDTA. Units of AP activity were calculated as described in [70].

### Dilution spotting

Primary cultures were pelleted, washed, and re-suspended in M9 minimal medium. Several dilutions of cultures were spotted on TBK-Brij and TBK-Ole plates supplemented with different concentrations of DTT. Plates were incubated and imaged (Gel Doc XR^+^ imaging system, Bio-Rad), at different time intervals. A representative image is shown in the figure.

### Determination of the redox state of DsbA protein

The redox state of DsbA was determined as described in [71], with slight modifications. Cells were treated with trichloroacetic acid (final concentration 20%) to prevent aerial oxidation and trap thiols of DsbA in their original state. Protein precipitates were collected by centrifugation, washed with acetone, dried, and dissolved in a freshly prepared solution containing 100 mM Tris-Cl (pH 8.8), 10 mM EDTA, 1% SDS and 30 mM AMS. Samples were incubated for 1 hour at 37°C in a thermomixer set at 1400 revolutions/min. Proteins were separated on a 15% non-reducing SDS-PAGE, transferred to nitrocellulose membrane and processed for Western blotting.

### Western Blotting

The redox state of DsbA and expression of AP was monitored by Western blot analysis. Samples were separated on SDS-PAGE and transferred to nitrocellulose membrane. The membrane was blocked with 5% (w/v) skimmed milk overnight at 4°C and probed with anti-DsbA (1:1000, Thermo Fisher Scientific) or anti-AP (1:5000, Millipore) primary antibody and HRP-conjugated anti-mouse (1:5000, Sigma) secondary antibody. Blots were developed using the SuperSignal West Dura Extended Duration Substrate (Pierce). Signal was captured on an X-ray film.

## Acknowledgments

We thank Abhijit Sardesai, Carol Gross, Christopher Rao, Manjula Reddy, and Sarah Ades for strains and plasmids. We thank members of the Chaba lab for discussions and critical reading of the manuscript.

## Supporting Information

### Supplementary Figure captions

**S1 Fig. LCFA-utilizing cells exhibit cadmium sensitivity.**

**S2 Fig. DsbA accumulates in a reduced form in WT MG1655 grown in LCFAs.**

**S3 Fig. Cpx pathway is considerably induced in the presence of DTT.**

**S4 Fig. NlpE is not the signal for Cpx induction during LCFA metabolism.**

### Supplementary Tables

**S1 Table. Strains and plasmids used in this study.**

**S2 Table. Primers used in this study.**

**Supplementary References**

