## Supplemental material for "Metabolism of long-chain fatty acids affects disulfide bond formation in *Escherichia coli* and activates envelope stress response pathways as a combat strategy"

### Supporting Information

#### Supplementary Figure captions

**S1 Fig. LCFA-utilizing cells exhibit cadmium sensitivity.** WT grown either in TBK-Brij or TBK-Ole and Δ*dsbA* grown in TBK-Brij were treated with different concentrations of $CdCl_2$ at time point T3. $OD_{600}$ of the cultures was measured after 24 hours of growth. Percent change in $OD_{600}$ of the cultures treated with $CdCl_2$ in comparison to their untreated controls was calculated and plotted against $CdCl_2$ concentration. Data represent the average (± S.D.) of three independent experiments. *Inset:* Growth curves of WT in TBK-Brij or TBK-Ole and Δ*dsbA* in TBK-Brij either without $CdCl_2$ treatment (open symbols) or treated with 2 mM $CdCl_2$ (filled symbols), is shown. Arrow indicates the time point T3 at which $CdCl_2$ was added. Data is shown for one of the three biological replicates.

**S2 Fig. DsbA accumulates in a reduced form in WT MG1655 grown in LCFAs.** WT MG1655 was grown either in TBK-Brij or TBK-Ole. Cultures were harvested at different time points as indicated in Fig. 5A, and processed as mentioned in the legend to Fig 4A. Δ*dsbA* and Δ*dsbB* cultured in TBK-Brij served as controls. Arrows indicate oxidized and reduced forms of DsbA**.**

**S3 Fig. Cpx pathway is considerably induced in the presence of DTT.** WT carrying either *cpxP-lacZ* or *rpoH*P3-*lacZ* reporter fusion was grown in TBK-Brij supplemented with increasing concentrations of DTT, as indicated. Cultures were harvested in the exponential phase, and β-gal activity was measured. Data were normalized to the β-gal activity of WT in TBK-Brij without DTT and represent average (± S.D.) of three independent experiments. The average β-gal activity of the *cpxP-lacZ* reporter strain in TBK-Brij without DTT was 13 (± 5) Miller units and that of *rpoH*P3-*lacZ* was 45 (± 17) Miller units.

**S4 Fig. NlpE is not the signal for Cpx induction during LCFA metabolism.** WT and Δ*nlpE* strains carrying *cpxP-lacZ* reporter fusion were grown either in TBK-Brij or TBK-Ole. Cultures were harvested in the stationary phase, and β-gal activity was measured. Data were normalized to the β-gal activity of WT in TBK-Brij and represent average (± S.D.) of three independent experiments. The average β-gal activity of the WT *cpxP-lacZ* reporter strain in TBK-Brij was 28 (± 5) Miller units.

### Supplementary Tables

**S1 Table. Strains and plasmids used in this study**

| Strain/Plasmid | Relevant genotype | Source [reference] |
|---|---|---|
| **Strains** | | |
| BW25113 | $F^-$ Δ(*araD-araB*)*567* Δ*lacZ4787*(::rrnB-3) $\lambda^-$ *rph-1* Δ(*rhaD-rhaB*)*568 hsdR514* | *E. coli* Genetic Stock Center |
| BW25142 | *$lacI^q$ rrnB3* Δ*lacZ4787 hsdR514* Δ(*araBAD*)*567* Δ(*rhaBAD*)*568* Δ*phoBR580 rph-1 galU95* Δ*endA9 uidA*(Δ*MluI*)::*pir-116 recA1* | Rao lab [1] |
| MG1655 | $F^-$ $\lambda^-$ *rph-1* | *E. coli* Genetic Stock Center |
| MG1655 Δ*lac* | MG1655 Δ*lacX74* | Gross lab |
| AM1247 | MG1655 *lacIZYA*::*frt nadA*::*Tn10 att*λ [$P_{pspA}$-*lacZ*], $Tet^r$ | Reddy Lab [2] |
| CAG45114 | MG1655 Δ*lacX74* λ[$P_{rpoHP3}$-*lacZ*] | Gross Lab [3] |
| CAG53524 | MC1061 [ϕλ*rpoH*P3::*lacZ*] *degS*ΔPDZ::*kan*, $Kan^r$ | Gross lab [4] |
| CAG55907 | CAG45114 Δ*relA* Δ*spoT*::*cam*, $Cam^r$ | Gross lab [5] |
| DH300 | MG1655 Δ(*argF-lac*)*U169* [$P_{rprA142}$-*lacZ*] | Ades Lab [6] |
| RI89 | $F^-$ Δ*araBAD714* (*araD139*)$_{B/r}$ Δ(*codB-lacI*)*3 phoR82 galK16 galE15* $\lambda^-$ *$e14^-$ relA1 rpsL150 spoT1 mcrB1* | Sardesai lab [7] |
| SEA4166 | MG1655 Δ*lacX74* λ*RS88* [$P_{cpxP}$-*lacZ*] | Ades Lab |
| SEA4170 | MG1655 Δ*lacX74* λ*RS88* [$P_{spy}$-*lacZ*] | Ades Lab |
| BW25113 Δ*fadL* | BW25113 *fadL::kan*, $Kan^r$ | Keio collection [8] |

| | | |
|---|---|---|
| BW25113 Δ*nuoK* | BW25113 *nuoK::kan*, Kan$^r$ | Keio collection [8] |
| BW25113 Δ*sdhB* | BW25113 *sdhB::kan*, Kan$^r$ | Keio collection [8] |
| BW25113 Δ*dsbA* | BW25113 *dsbA::kan*, Kan$^r$ | Keio collection [8] |
| BW25113 Δ*dsbB* | BW25113 *dsbB::kan*, Kan$^r$ | Keio collection [8] |
| RC15082 | BW25113 *att*λ::[Kan $P_{fadE}$ -*lacZ oriR6K*], Kan$^r$ | This work |
| RC5264 | P1 (BW25113 *fadL*::*kan*) X RI89, Kan$^r$ | This work |
| RC15062 | P1 (BW25113 *fadE*::*kan*) X RI89, Kan$^r$ | This work |
| RC5327 | P1 (BW25113 *dsbA*::*kan*) X RI89, Kan$^r$ | This work |
| RC5328 | P1 (BW25113 *dsbB*::*kan*) X RI89, Kan$^r$ | This work |
| RC5325 | P1 (BW25113 *cpxR*::*kan*) X RI89, Kan$^r$ | This work |
| RC5333 | RC5325, *kan* cassette flipped out | This work |
| RC5326 | P1 (CAG53524) X RI89, Kan$^r$ | This work |
| RC5337 | P1 (BW25113 *dsbA*::*kan*) X MG1655, Kan$^r$ | This work |
| RC5338 | P1 (BW25113 *dsbB*::*kan*) X MG1655, Kan$^r$ | This work |
| RC15106 | P1 (AM1247) X MG1655 Δ*lacX74* $nadA^+$ | This work |
| RC15006 | P1 (BW25113 *fadL*::*kan*) X CAG45114, Kan$^r$ | This work |
| RC15005 | P1 (BW25113 *fadE*::*kan*) X CAG45114, Kan$^r$ | This work |
| RC5317 | P1 (CAG53524) X CAG45114, Kan$^r$ | This work |
| RC15104 | P1 (BW25113 *rseB*::*kan*) X CAG45114, Kan$^r$ | This work |
| RC15103 | RC15104, *kan* cassette flipped out | This work |
| RC15105 | P1 (BW25113 *relA*::*kan*) X CAG45114, Kan$^r$ | This work |
| RC15095 | P1 (BW25113 *cydD*::*kan*) X CAG45114, Kan$^r$ | This work |
| RC15020 | P1 (BW25113 *fadL*::*kan*) X SEA4166, Kan$^r$ | This work |
| RC15019 | P1 (BW25113 *fadE*::*kan*) X SEA4166, Kan$^r$ | This work |
| RC15043 | P1 (BW25113 *ackA*::*kan*) X SEA4166, Kan$^r$ | This work |
| RC5320 | RC15043, *kan* cassette flipped out | This work |
| RC5322 | P1 (BW25113 *pta*::*kan*) X RC5320, Kan$^r$ | This work |
| RC5302 | P1 (BW25113 *cpxA*::*kan*) X SEA4166, Kan$^r$ | This work |
| RC15096 | P1 (BW25113 *cydD*::*kan*) X SEA4166, Kan$^r$ | This work |

| | | |
|---|---|---|
| RC18029 | P1 (BW25113 *nlpE*::*kan*) X SEA4166, Kan$^r$ | This work |
| **Plasmids** | | |
| pCP20 | pSC101 *ori cI857* $\lambda$-$P_R$ *flp* ts Amp$^r$ Cam$^r$ | [9] |
| pINT-ts | *oriR6K int* Amp$^r$ | Rao lab [1] |
| pAH125 | *oriR6K,* MCS-*lacZ t0 att*$\lambda$ Kan$^r$ | Rao lab [1] |
| pMS02 | *oriR6K,* MCS $P_{fadE}$-*lacZ t0 att*$\lambda$ Kan$^r$ | This work |

#### S2 Table. Primers used in this study

| Primer Name | Purpose | Sequence (5'-3') |
|---|---|---|
| **Primers used for cloning *fadE* cis element in pAH125 and its verification** | | |
| MS43 | Forward primer for cloning *fadE* promoter in pAH125 | ACCGGGTACCATTGATTTAAGAATTTTCAGGTCGGATGC |
| MS44 | Reverse primer for cloning *fadE* promoter in pAH125 | CGTGAATTCCCGTAGCGAGAATACTCAAAATCATCAT |
| BS106 | Sequencing/verification primer for cloning in pAH125 | TTGTCGGTGAACGCTCTCCT [10] |
| MS49 | Sequencing/verification primer for cloning in pAH125 | TAAAACGACGGCCAGTGAATCC |
| Restriction sites are underlined | | |
| **Primers used for strain verification of single integrants at the *att*$\lambda$ site** | | |
| GA22 | Forward primer specific to *att*$\lambda$ site on *E. coli* chromosome for confirmation of single-copy integration of reporter plasmids | GGCATCACGGCAATATAC [10] |
| GA23 | Forward primer specific to pAH125 to identify multiple-copy integration of reporter plasmids | ACTTAACGGCTGACATGG [10] |
| GA25 | Reverse primer specific to *att*$\lambda$ site on *E. coli* chromosome for confirmation of single-copy integration of reporter plasmids | TCTGGTCTGGTAGCAATG [10] |
| GA29 | Reverse primer specific to pAH125 to identify multiple-copy integration of reporter plasmids | TGCGAGGCTTTGTGCTTC [10] |

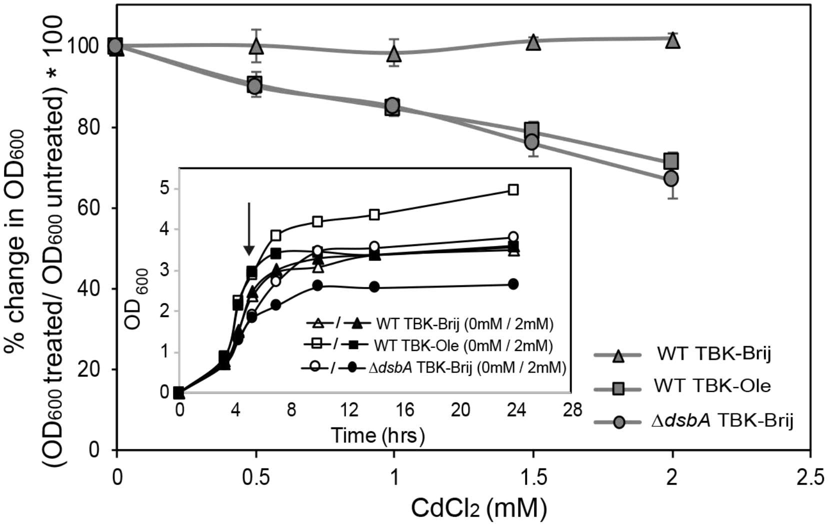

% change in OD600
(OD600 treated/ OD600 untreated) * 100
0
20
40
60
80
100
0
0.5
1
1.5
2
2.5
CdCl2 (mM)
WT TBK-Brij
WT TBK-Ole
ΔdsbA TBK-Brij
OD600
0
1
2
3
4
5
0
4
8
12
16
20
24
28
Time (hrs)
WT TBK-Brij (0mM / 2mM)
WT TBK-Ole (0mM / 2mM)
ΔdsbA TBK-Brij (0mM / 2mM)


Figure S1

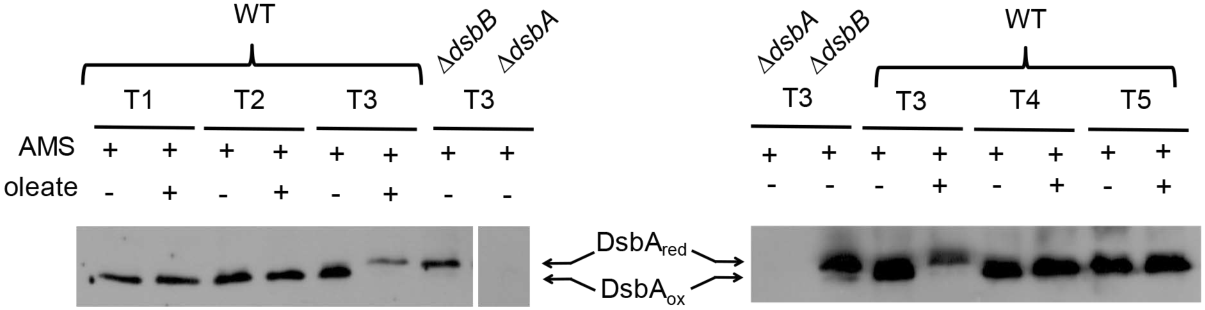
WT
ΔdsbB
ΔdsbA
T1
T2
T3
T3
AMS
oleate
DsbA_red
DsbA_ox
ΔdsbA
ΔdsbB
WT
T3
T3
T4
T5

Figure S2

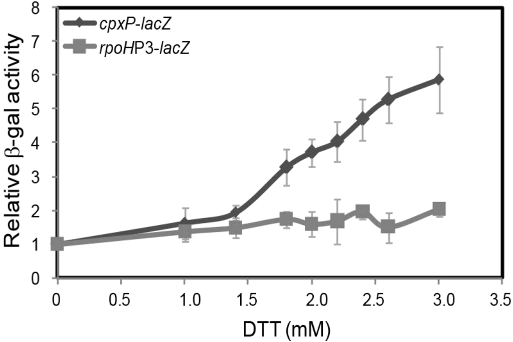

cpxP-lacZ
rpoHP3-lacZ
Relative β-gal activity
0
1
2
3
4
5
6
7
8
0
0.5
1.0
1.5
2.0
2.5
3.0
3.5
DTT (mM)


### Figure S3

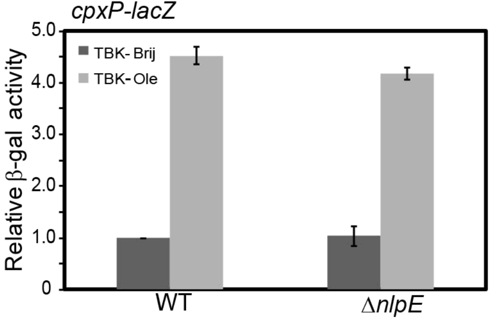
cpxP-lacZ
5.0
4.0
3.0
2.0
1.0
0
Relative β-gal activity
TBK- Brij
TBK- Ole
WT
ΔnlpE

Figure S4
